# Whole Slide Preview Image Segmentation and Setup for Digital Pathology Scanners

**DOI:** 10.1101/2020.02.24.963645

**Authors:** Mahdi S. Hosseini, Dohyoung Lee, Daniel Gershanik, Dongwoon Lee, Savvas Damaskinos, Konstantinos N. Plataniotis

## Abstract

The problem of tissue finding is of special interest in automating WSI scanners where it decomposes the preview image of tissue glass slides into a simplified and abstract level of localization and identification to setup WSI scanner for high-resolution scan. Prior to such scanning, a preview image is captured to calibrate the scanner’s parameters. Scan parameters such as focus depth and scan region are determined using a tissue finding software package. This paper introduces a series of pipelines (e.g. binary mask segmentation, tissue/artifact classification, region-of-interest allocation) to automate tissue preview segmentation in both brightfield and darkfield microscopy.

## I. Introduction

Digital pathology (DP), which also known as “virtual microscopy” or “whole slide imaging (WSI)”, refers to the process of visualizing, navigating, focusing, marking, and classifying the areas of interest within the glass tissue slides on a computer monitor [1]. DP is a computer oriented technology environment that facilitates the digitalization of images from biological tissues, collected on glass slides, by a computing device integrated with an optical scanning device. It is considered as an alternative technology to the traditional optical microscopy undertaken by a skilled professional for imaging biological tissues, and it can enable a completely digital workflow for anatomical pathology departments. The demand for such fully digital processing workflow has increased exponentially mostly due to its use in monitoring and primary diagnosis from disease tissues of human specimen [1]–[4].

Routine diagnosis of such pathology slides requires high quality high throughput images. High quality images from the slides must be displayed in a consistent and reliable manner to assist research pathologists to interpret these images accurately and efficiently. It is noteworthy that the US Food and Drug Administration (FDA) provides a guideline that requires vendors of digital pathology (DP) scanners to integrate viable image management software to include tissue finding, focus mapping, optical correction, and color sensor calibration for image reproducibility [5], [6]. Despite effective utilization of existing virtual microscopes, the goal of delivering a fully integrated, scalable, certified for clinical use, digital pathology solution has not yet materialized. According to early adopters of integrated digital pathology the main reason for the delay is the lack of progress in integrating scanners with the image analysis modules. The delay could be attributed to issues related to: a) Hardware-end configuration: This processing pipeline comprises of several function blocks designed to automate the image acquisition process such as tissue allocation, focus detection, stitching and storage of high-resolution images, and b) Software-end configuration: Each manufacture utilizes a proprietary protocol for the image software algorithms utilized in the pipeline resulting in a diverse set of image navigation, contrast correction, color enhancement, segmentation, and information retrieval solutions. The overall scanner-guided image analysis software is demonstrated in Figure 1.

**Fig. 1.**
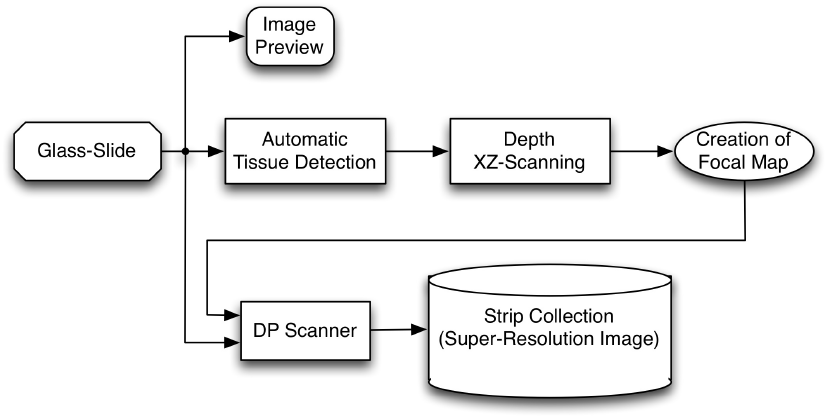
Whole Slide Imaging (WSI) scanner-guided image analysis software. The glass-slide is previewed and region-of-interests of tissues are identified, focus-dots are allocated on the tissue areas and focus-map is created to guide the high-resolution tissue scan where image strips are collected at the end.

The tissue finding package segments the preview image, classifies the preview image, places focus dots and a white balance box in order to guide the WSI scanners through high optics magnification scans. The same software package is also needed for darkfield imaging. Fluorescence microscopy uses a phosphorfluorescence dye that is applied to tissue samples to highlight specific regions and properties within the tissue sample. Because phosphorfluorescence dye is clear, fluorescence tissue samples are not visible in brightfield preview images. Hence, darkfield illumination must be used to take preview images. Darkfield illumination is a microscopy method that excludes direct light, only including scattered light in the image.

The purpose of this study is to develop a user-friendly Tissue Finding Package (TFP) software in order to segment, classify, place focus-dots and white balance box for both brightfield and darkfield preview images. The TFP software is to automatically guide WSI scanner throughout high resolution scan.

## II. Brightfield Tissue Finding Package

The brightfield preview segmentation package is developed by the University of Toronto’s Multimedia lab and was subsequently ported to C++ by Huron’s technical team. The segmentation pipeline is described in detail in figure 2. The segmentation package consists of a four-step pipeline from beginning to end. The four major steps, initialization, segmentation, classification and ROI analysis are comprised of sub steps as seen in figure 2. Initialization is the first step in the segmentation pipeline. It consists of two sub steps: initialize parameters and scan tissue for preview imaging of tissue glass slides. Initialize parameters is the first step in the tissue segmentation pipeline that sets up the scanner for preview imaging. Once the preview image is obtained, it is fed into segmentation pipeline to segment potential tissue regions. The tissue segments are fed to classification pipeline to discriminate between real tissues and possible artifacts. The final tissue regions are then provided to user/scanning system where regions-of-interest (ROI) are identified for high-resolution tissue scans.

**Fig. 2.**
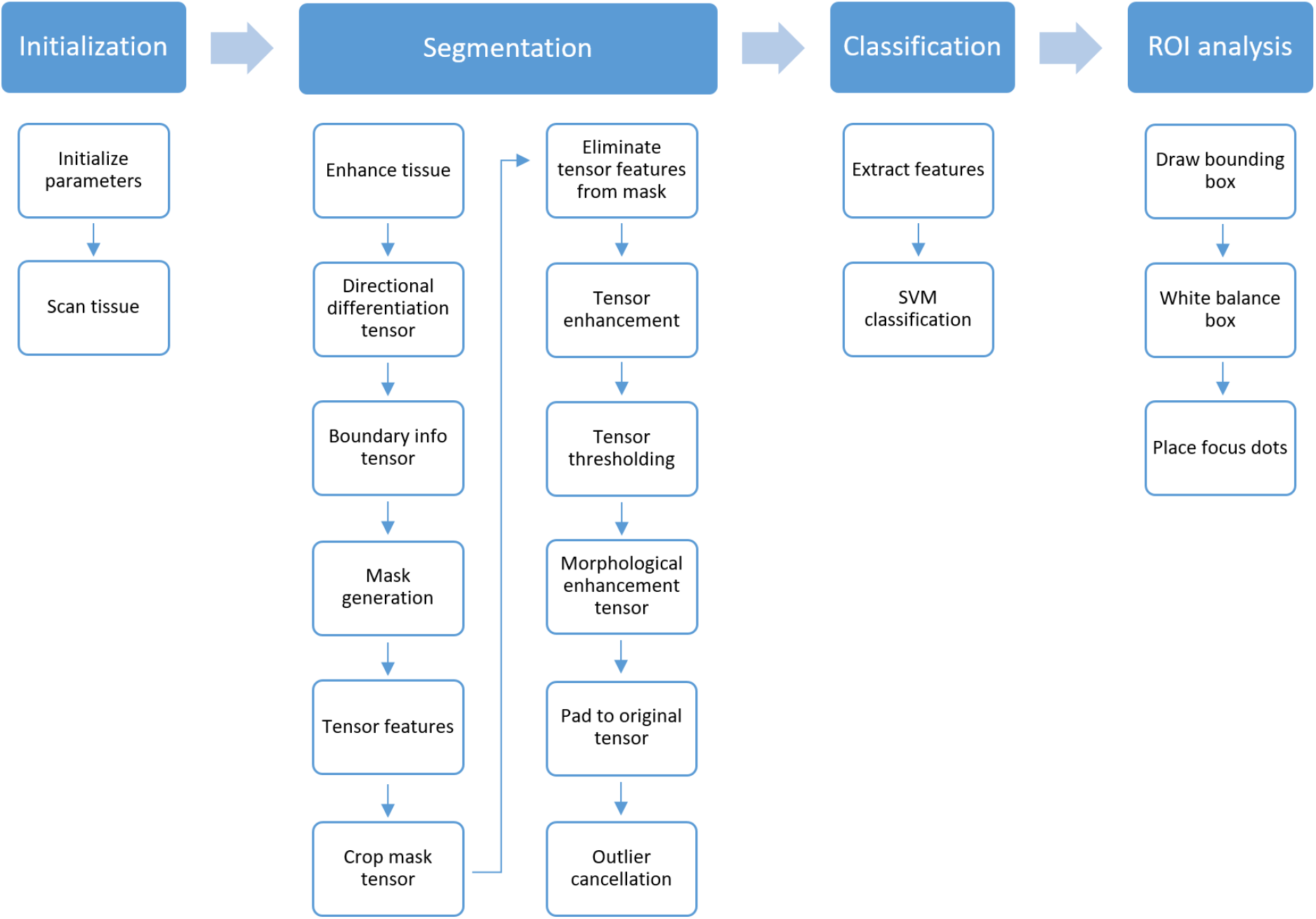
Brightfield tissue segmentation pipeline. The pipeline consists of four major steps: initialization, segmentation, classification and ROI analysis. Each major step consists of sub-steps, drawn in blue outlined boxes. The initialization step defines constants and initializes global variables used throughout the pipeline. The segmentation step identifies objects in the image and separates them for the background. The classification step determines which objects in an image are tissue samples and which are not. ROI analysis step callibrates the high magnification image capture parameters.

## III. Segmentation

The segmentation step separates the image’s features from its background. The pipeline utilizes both threshold masking and a Frangi filter [7] in parallel to achieve optimal segmentation results. Note that the derivative operators of Frangi filter are replaced by MaxPol library package introduced in [8], [9] for high accuracy approximation. The mask and Frangi filter outputs are then superimposed at the Eliminate tensor features from mask step. This process is illustrated in figure 3.

**Fig. 3.**
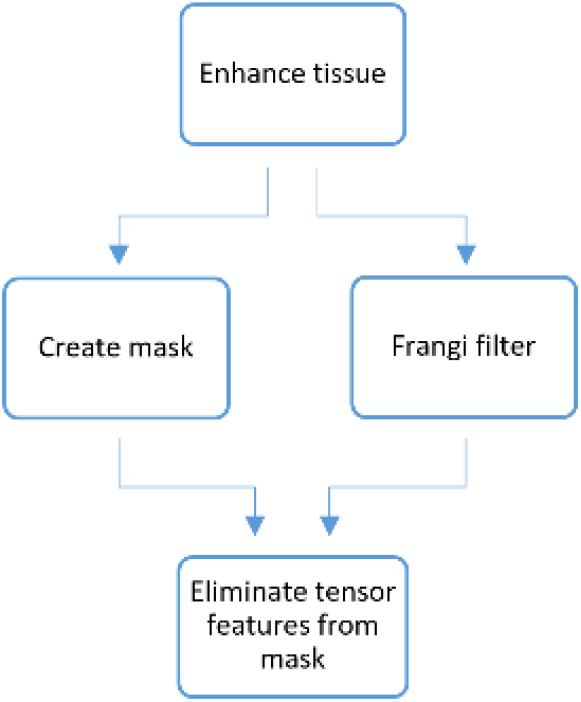
Frangi filter and masking steps both use the enhanced image as input. The outputs are then combined in the Eliminate tensor features from mask step.

### A. Tissue Enhancement

The first sub step is tissue enhancement, here the image is first denoised using pixel averaging. Subsequently, colour correction based on background average is preformed. This denoised image is used as an input for both the Frangi Filter and the threshold masking steps. LAB and HSV color versions of the image are also created here. Figure 4 shows the outputs from this sub-step.

**Fig. 4.**
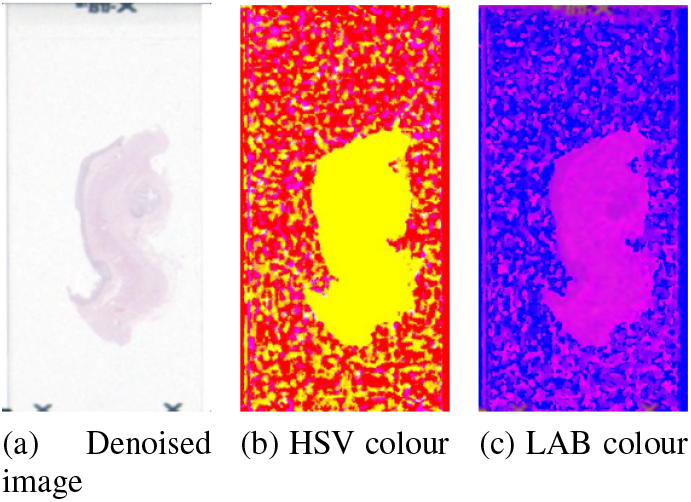
Tissue enhancement step produces takes the original images as input and produces three output images. A denoised image in RGB a denoised image in HSV colours and a denoised image in LAB colours.

### B. RGB and L channel thresholder mask

Directional differentiation tensor, boundary info tensor, mask generation and crop mask tensor are sub steps used to create the threshold mask. The directional differentiation sub step creates first and second derivatives of the RGB and LAB L channel images. These are used in the boundary info substep to find the crop boundaries for the original image. The mask generation step creates an RGB mask and a LAB L channel mask using an adaptive threshold. These two masks are then superimposed using an element wise AND function. Finally the mask edges are cropped using the previously determined crop boundaries. The threshold used is designed to be forgiving so that all of the tissue is found on the mask in addition to artifacts. Figure 5 shows the RGB mask, L mask and the combined mask.

**Fig. 5.**
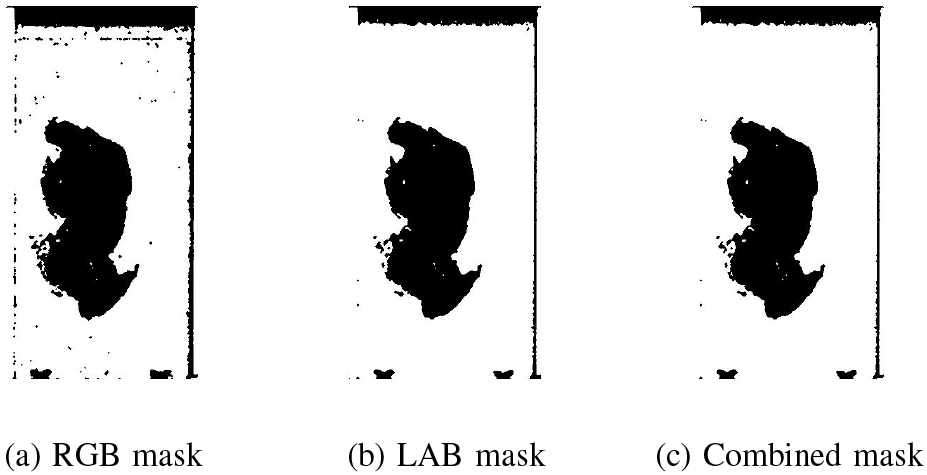
Mask generation step produces two masks, an RGB mask by thresholding the RGB denoised image and a LAB L channel mask by thresholding the LAB L channel image. The two masks are then combined using a AND function.

### C. Tensor features

As previously mentioned, in addition to threshold masking the pipeline uses a Frangi filter for further image segmentation. The denoised image is passed through a Frangi filter, which highlights ridges and vesicles of the image, seen in figure 6 a.

**Fig. 6.**
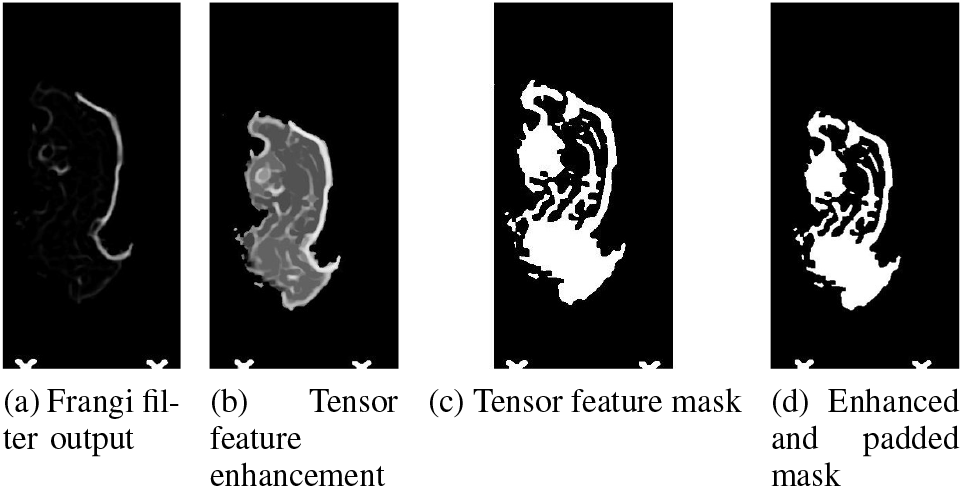
Frangi filter output manipulation. All tensor features not found on the thresholded masks are removed and then enhanced using a log function. The output is then thresholded, padded to original size and enhanced using morphological enhancement.

#### 1) Eliminate tensor features from mask

The next step, eliminate tensor features from mask removes all the features not found in the mask from the Frangi filter output found in the previous step. This is done to remove all the features from the Frangi output that are certainly not tissue while maintaing feature that could be tissue for further segmentation. The output from this step is shown in figure 6.

### D. Tensor enhancement and morphological enhancement

The tensor enhancement step highlights the Frangi’s output features, seen in figure 6 b. A log function is used to separate low and high strength signals. Further, morphological enhancement is used to fill in the holes created between the ridges and vesicles outputted by the Frangi filter, seen in figure 6 d.

### E. Tensor thresholding

The Frangi output is then thersholded to create a new mask. The new mask incorporates the vesicle information extracted from the Frangi filter, combined with the feature elimination from the previous mask. It therefore produces finer segmentation than the original RGB and L channel masks. Once the mask is produces it undergoes morphological enhancement.

### F. Pad to original tensor

In the pad to original tensor step the previously cropped areas are restored by padding zeros and removing small artifacts, which are too small to be tissue.

### G. Cancel outliers

The final step prior to image classification is outlier cancellation. Here all regions on the segmented map below a certain size are removed, since they are too small to possibly be tissue.

## IV. Feature Extraction from Tissue Segments

The segmentation process discussed in the previous section generates a binary spatial map of potential tissue regions (i.e., clusters) for an input digital slide image. However, the segmented map may still contain non-tissue objects due to the diverse artifacts present in a slide image and the limitations in segmentation. Therefore, non-tissue objects detected as tissues should be selectively eliminated from a binary map while preserving true tissue objects. This process can be formulated as a binary object classification problem of tissue and nontissue objects.

Initially, individual connected regions should be identified from a binary map by connected component labelling to facilitate object-level analysis. Connected component labelling works by scanning a binary image from segmentation process and assigning unique labels to connected pixel regions, i.e. regions of adjacent pixels which share the same set of intensity values *V* (For a binary image, *V* = 1). Therefore, it is highly dependent on underlying segmentation mechanism. In this work, we assume 8-connectivity for connected component labelling. Fig. 7 exemplifies the results of connected component labelling on a sample H&E image containing more than 30 connected regions. Subsequently, individual connected regions from the segmented images should be classified into either tissue and non-tissue regions.

**Fig. 7.**
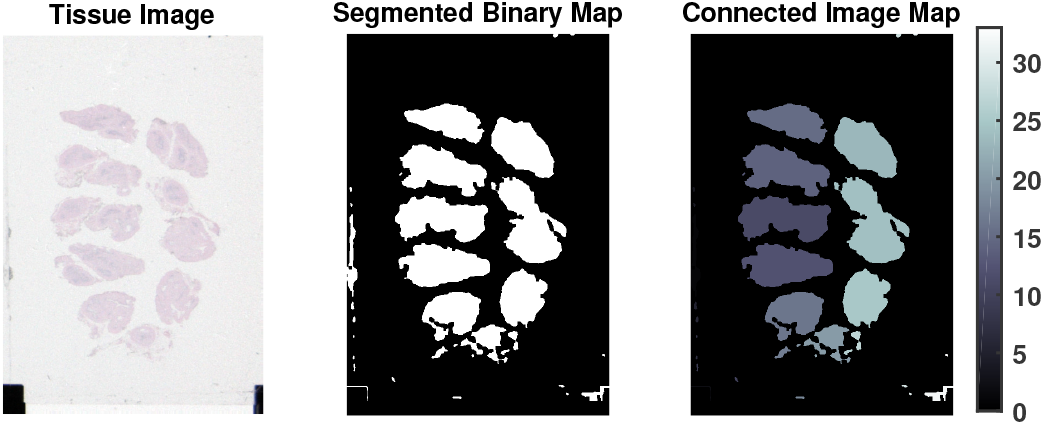
Example of connected component labelling on a sample H&E image. individual connected regions are indicated by different levels of gray intensity.

The extraction of image features is a crucial step for tissue classification task. Desirable image features for classification convey discriminative information of tissue objects while do not require excessive computational costs. Since it is practically impossible to find a single universal image feature that works perfectly on diverse tissue types shown in Table, we consider a combination of low-level image features that are believed to be related to the classification of tissue and non-tissue clusters. The features selected for our analysis can be categorized into several classes: i) color (intensity); ii) texture; and iii) geometric. The complete set of exploited image features and their descriptions are provided in Table III. compared with the existing works on tissue classification, our work has following contributions:

- We advance the research field by demonstrating the effectiveness of commonly used low-level image features for the classification of tissue objects. The image statistics provided in this work can be used as the guideline for the preview image analysis in whole slide imaging (WSI) technology.
- The hue-based image features are properly extracted using directional statistics considering the periodicity of hue information. In addition, they are weighted by pixelwise saturation information in order to take into account the reliability of hue information.

**TABLE I.** Descriptions of the manually labelled tissue and non-tissue datasets

| Data Set | No. Image | Image Format | No. Tissue/Nontissue | Remarks |
| --- | --- | --- | --- | --- |
| Mixed Previews | 124 | TIFF, 260×530<br>468×680 | 781 / 866 | High to mid quality images with white or grayish background |
| Tissue Snap Preview | 120 | JPEG, 273×530 | 610 / 790 | Low quality (blurred) images with red casting issue |
| Set 1 | 150 | TIFF, 272×555 | 885 / 681 | High quality images with bright white background |
| **Total** | **394** |  | **2276 / 2337** |  |

**Table II.** Selected Features for Different Selection Techniques

| Feature # | 1 | 2 | 3 | 4 | 5 | 6 | 7 | 8 | 9 | 10 | 11 | 12 | 13 | 14 | 15 | 16 | 17 | 18 | 19 | 20 | 21 | 22 | 23 | 24 | 25 | 26 |
| --- | --- | --- | --- | --- | --- | --- | --- | --- | --- | --- | --- | --- | --- | --- | --- | --- | --- | --- | --- | --- | --- | --- | --- | --- | --- | --- |
| Fisher | 28 | 27 | 11 | 2 | 12 | 17 | 24 | 16 | 4 | 25 | 7 | 21 | 15 | 19 | 18 | 10 | 6 | 20 | 26 | 8 | 23 | 5 | 3 | 9 | 13 | 14 |
| mRMR-MI(D) | 28 | 2 | 27 | 12 | 21 | 9 | 11 | 23 | 14 | 6 | 17 | 8 | 16 | 3 | 25 | 13 | 18 | 24 | 7 | 19 | 4 | 26 | 20 | 5 | 15 | 10 |
| mRMR-MI(Q) | 28 | 27 | 12 | 6 | 16 | 3 | 11 | 21 | 7 | 17 | 19 | 2 | 25 | 5 | 24 | 14 | 18 | 15 | 23 | 9 | 10 | 4 | 26 | 8 | 20 | 13 |
| mRMR-FC(D) | 28 | 17 | 27 | 14 | 2 | 16 | 3 | 21 | 11 | 25 | 4 | 8 | 24 | 12 | 20 | 9 | 18 | 26 | 7 | 13 | 23 | 19 | 6 | 15 | 10 | 5 |
| mRMR-FC(Q) | 28 | 17 | 27 | 2 | 16 | 11 | 24 | 21 | 12 | 25 | 4 | 7 | 8 | 20 | 18 | 26 | 19 | 15 | 6 | 10 | 3 | 23 | 9 | 5 | 13 | 14 |

**Table III.**
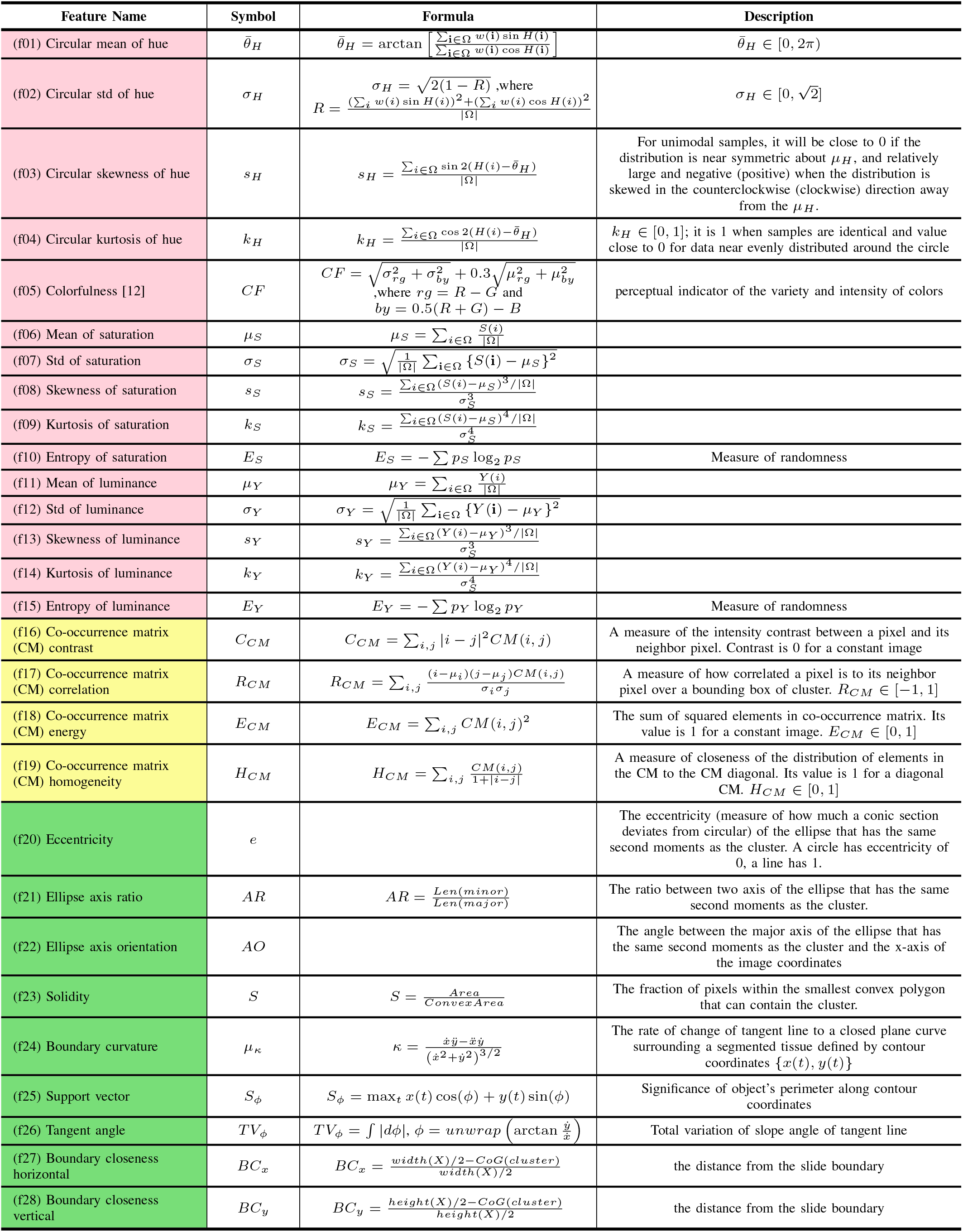
List of the image features used for the classification of tissue and non-tissue clusters.

The workflow of the feature extraction process in this section is demonstrated in Fig. 8.

**Fig. 8.**
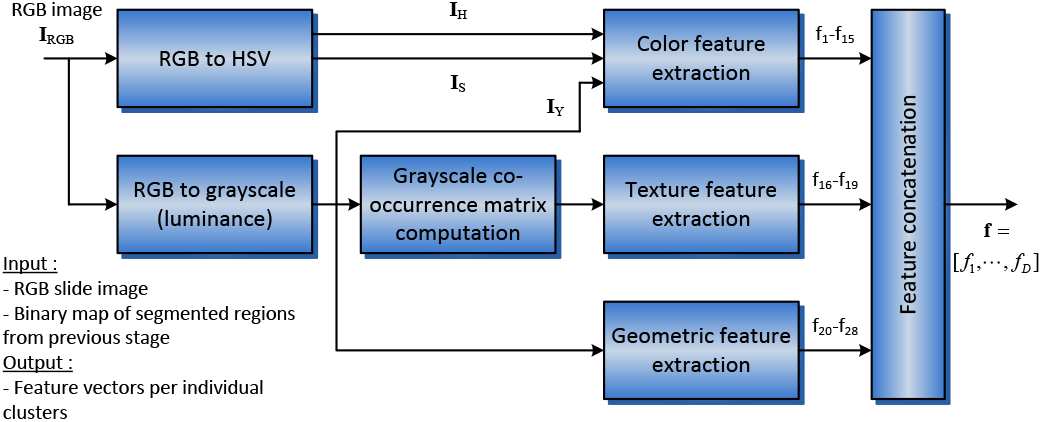
Illustration of the feature extraction pipeline

### A. Color (intensity)-based Features

These features provide quantitative information about the pixels within a segmented region, extracted from the luminance (gray-level) or color channels of slide images. They do not provide any information about the spatial distribution of the pixels but provide rotation and scaling invariant information. The color features are extracted from following three color channels: hue, luminance and saturation.

#### 1) Hue-based Features

Hue is a perceptual attribute related to the dominant wavelength of a color signal, represented as angular quantity in cylindrical color spaces. Hue provides crucial information regarding the tissue type since varying tissue types exhibit different stain colors. The hue representation is obtained by transforming RGB input to HSV color space [10]. The hue-based features considered in this paper differs from the existing methods in following ways: i) hue information in preview tissue images are processed with directional statistics to properly take into account its angular nature,; ii) the saturation weighted scheme is used for the extraction of hue features in order to enhance the reliability of hue information. The use of saturation-weighted scheme is effective for collecting reliable hue statistics since it reduces the contribution of hue information from either achromatic (desaturated) tissue regions or white background regions (falsely included into potential cluster regions due to inaccurate segmentation).

##### Circular mean of hue

The circular mean of hue 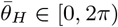 can be obtained by:

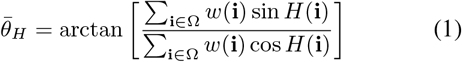

where the i ∈ ℝ^2^ is a spatial coordinate, the Ω is a set of pixel coordinates of potential tissue cluster and the operator |·| indicates the cardinality of a set. The spatial weighting factor {*w*(**i**)|**i** ∈ Ω} is introduced to assign more importance on highly saturated pixels since hue value of achromatic pixel (i.e., desaturated pixel) has no significance as small perturbations in pixel value may result in significant hue changes. In addition, if tissue objects contain some background pixels due to suboptimal segmentation process^1^, the proposed saturation-based weighting scheme can effectively reduce the contribution of background pixels in the computation of 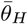 since background pixels typically have low saturation values. We exploit a sigmoid curve to define the weighting function of smooth transition between chromatic and achromatic pixels (Fig. 9):

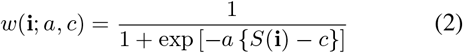

where the *c* determines the point for which the significance level is equal to 0.5 and the *a* is the curvature parameter (the smaller the *a*, the smoother the curve). In this work, two parameters are selected empirically selected as (*c,a*) = (0.2,40). Fig. 10 demonstrates that the saturation-weighted circular mean descriptor yields more accurate description of color information than the non-weighted descriptor.

**Fig. 9.**
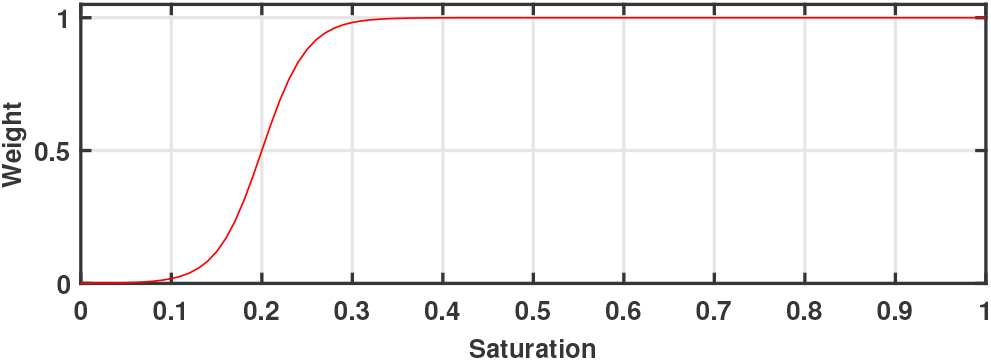
The sigmoid function used for describing the saturation-weighted scheme.

**Fig. 10.**
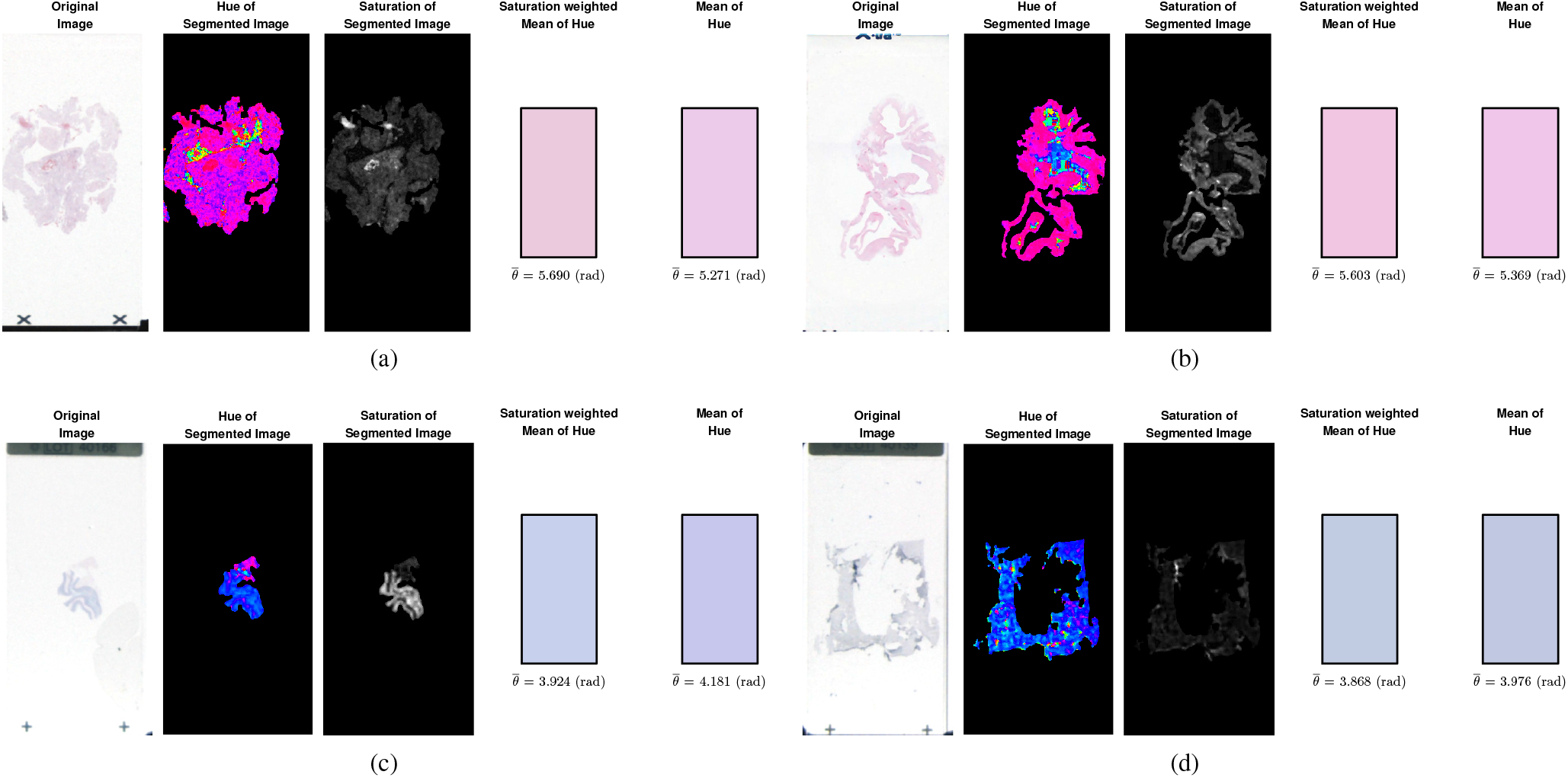
Comparison of the circular mean of hue with and without the saturation-weighted scheme. The second and third images demonstrates the hue and saturation channels of tissue regions.

##### Circular standard deviation of hue

The circular standard deviation of hue 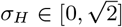 can be obtained by:

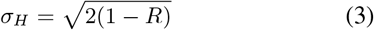

where the mean resultant vector *R* is given by:

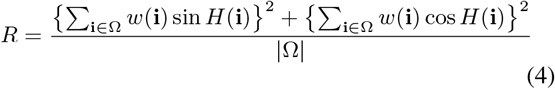

The closer the circular standard deviation to zero, the more concentrated the hue sample is around the circular mean.

##### Circular skewness of hue

The circular skewness measures the degree of symmetry in circular distribution [11]:

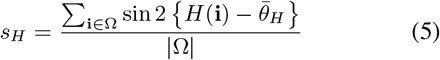

For unimodal samples, it will be close to zero if the distribution is near symmetric about 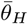 and relatively large and negative (positive) when the distribution of the data is skewed in the counterclockwise (clockwise) direction away from the 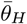.

##### Circular kurtosis of hue

The circular kirtosis measures the degree of peakedness in circular distribution [11]:

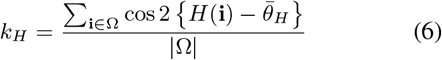

where it takes the maximum value of one when samples are identical and value close to zero for data near evenly distributed around the circle

#### 2) Luminance- and Saturation-based Features

In this work, we make use of first four moments of luminance and saturation channels to characterize objects. The saturation of color stimuli is related to the purity of color. The first four color moments for color channel *C* ∈ {*Y, S*} are given by:

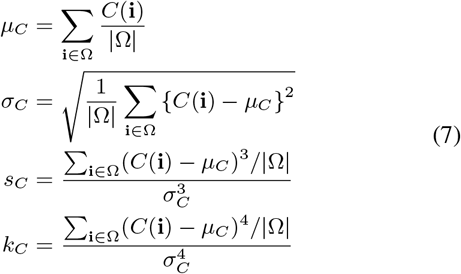

where *μ_C_, σ_C_, s_C_* and *k_C_* are the mean, standard deviation, skewness and kurtosis of color channel *C*.

In addition, the entropies of both saturation and luminance channels are computed. Given a color channel *C*, the channel entropy is defined by:

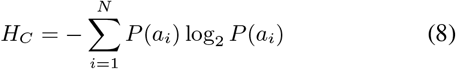

where *N* is the total number for intensity levels for color channel, *a_i_* is the *i*-th level and *P*(*a_i_*) is the probability of the *i*-th level (estimated from the image histogram)/

#### 3) Colorfulness Feature

Colorfulness is a perceptual indicator of the variety and intensity of colors, defined as follows [12]:

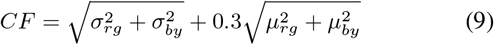

where *r_g_* and *y_b_* are the simple opponent color spaces defined as *r_g_* = *R – G* and *y_b_* = 0.5(R + *G*) – *B*, respectively.

### B. Geometric Features

The geometric features provide information about the size and shape of a region. The size is expressed by the radius, area, and perimeter of the cell. On the other hand, the shape is expressed by the compactness, roundness, smoothness, length of the major and minor axes, symmetry, concavity, and perimeter. For this section, we assume the binary map *B_l_*(*x, y*) has a value of 1 for the pixel coordinates (*x,y*) corresponds to *l*-th tissue region, otherwise has a value 0.

First three geometric features are obtained from the fitted ellipse with the same second moments as the potential tissue regions. Let 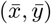 are the centroid of the object given by:

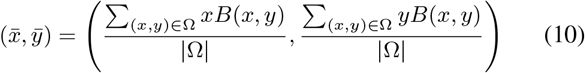

The lengths of major and minor axis for the fitted ellipse, denoted as *a,b*, are computed by:

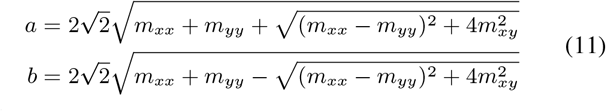

where

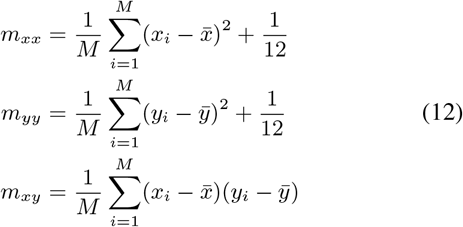

The eccentricity is a measure of how much a conic section deviates from circle, where a circle has eccentricity of 0, and a line has eccentricity of 1.

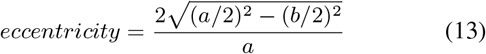

The ratio between the ellipse axis *AR* is defined as:

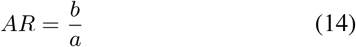

The ellipse axis orientation is defined as an angle between the major axis of the ellipse and the x-axis as follows:

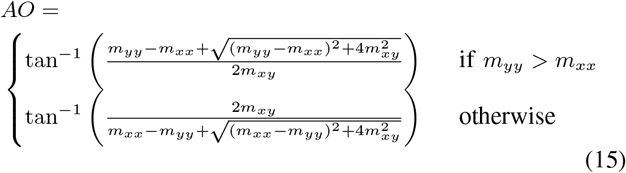

The solidity measures the fraction of pixels within the convex hull that are within the cluster region:

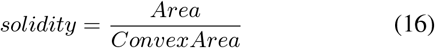

where the solidity of 1 indicates a solid object while the solidity less than 1 indicates irregular region or holes.

The horizontal and vertical boundary closeness, denoted as *BC_x_* and *BC_y_* provides the normalized distances from the centroid of tissue cluster to the slide boundary as follows:

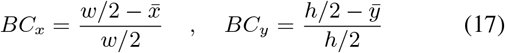

where *w* and *h* are the width and height of the slide image. They are meaningful to identify clusters since regions located near the boundary of slide are highly likely non-tissues.

### C. Texture Features

Image texture provides information about the variation in the intensity of a surface by quantifying properties such as smoothness, coarseness, and regularity. In order to extract textural features, the graylevel co-occurrence matrix (CM) is computed from the luminance channel since it is widely used to capture the spatial dependence of luminance values which contribute to the perception of texture.

The co-occurrence matrix describes the number of occurrences of gray-level *i* in a specific spatial relation with graylevel *j*. Spatial relations are typically specified by distance *d* and direction *θ*. The graylevel CM can be defined as:

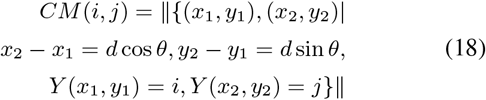

where (*x*_1_,*y*_1_) and (*x*_2_,*y*_2_) are pixel positions, *Y*(·) is the luminance value of the pixel and || · || is the number of the pixel pairs that satisfy the conditions. The number of luminance levels has an impact to the computational cost of the texture statistics. The quantization merges similar luminance levels within the image and reduces the influence of noise or artifacts [13]. For this project, the luminance intensity is discretized to 64 levels and the co-occurrence matrix is obtained for a distance of *d* = 1 and four directions *θ* = 0°, 45°, 90°, 135°. For the regions of tissue candidates with non-rectangle shape, the axis-aligned minimum bounding rectangle is used to compute the co-occurrence matrix. Assuming that textures in slide image does not have specific directions, four texture descriptors in (19) are obtained by averaging the co-occurrence matrices of all four directions.

The *contrast* measures the amount of local variations present in the image. A low value of contrast is obtained when the image has almost constant luminance values. The *correlation* measures how correlated a pixel is to its neighbor over the whole image. A value of 1 is obtained from an area with constant luminance value, indicating perfect correlation. The *energy* measures the texture uniformity. It is high when the GLCM has few entries of large magnitude, low when all entries are almost equal. The *homogeneity* measures the closeness of the distribution of elements in the GLCM to the GLCM diagonal. It produces high value for areas with little difference in luminance, which produces a GLCM with nonzero values concentrated along or near the main diagonal.

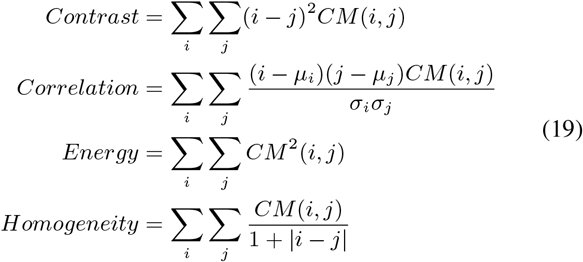

### D. Extracted Feature Analysis

Although we have carefully chosen aforementioned 28 image features due to their relevances to tissue object characterization, dealing with high dimensional data (i.e., data with large number of features) are not desirable for classification tasks due to the curse of dimensionality. In the presence of redundant, irrelevant or noisy features, classification methods become less interpretable and may lead to overfit. Therefore, removing such features leads to the reduction of data acquisition cost and training delay, the enhancement of learning accuracy and the improvement of classifier comprehensibility. In this section, we evaluate the effectiveness of individual features in terms of their *relevance* and *redundancy*. Here, the former indicates the relevance of features to the labelled class while the latter indicates the redundancy between features. Ideally, one should select image features that are highly relevant to the labelled classes while avoiding the redundancy which may appears when two relevant features are closely associated to each other.

Fig. 11 demonstrates the histograms of individual image features for labelled tissue and non-tissue samples. The distribution patterns reveals that individual features have varying degrees of class relevance (i.e., separability). For instance, the higher order color statistical features (e.g., the skewness and kurtosis of luminance and saturation channels) provide insignificant discriminative powers as the histograms for both classes substantially overlap. On the other hand, some geometric features, in particular, two boundary closeness features provides strong discriminative powers as they have two distinct peaks for tissue and non-tissue classes.

**Fig. 11.**
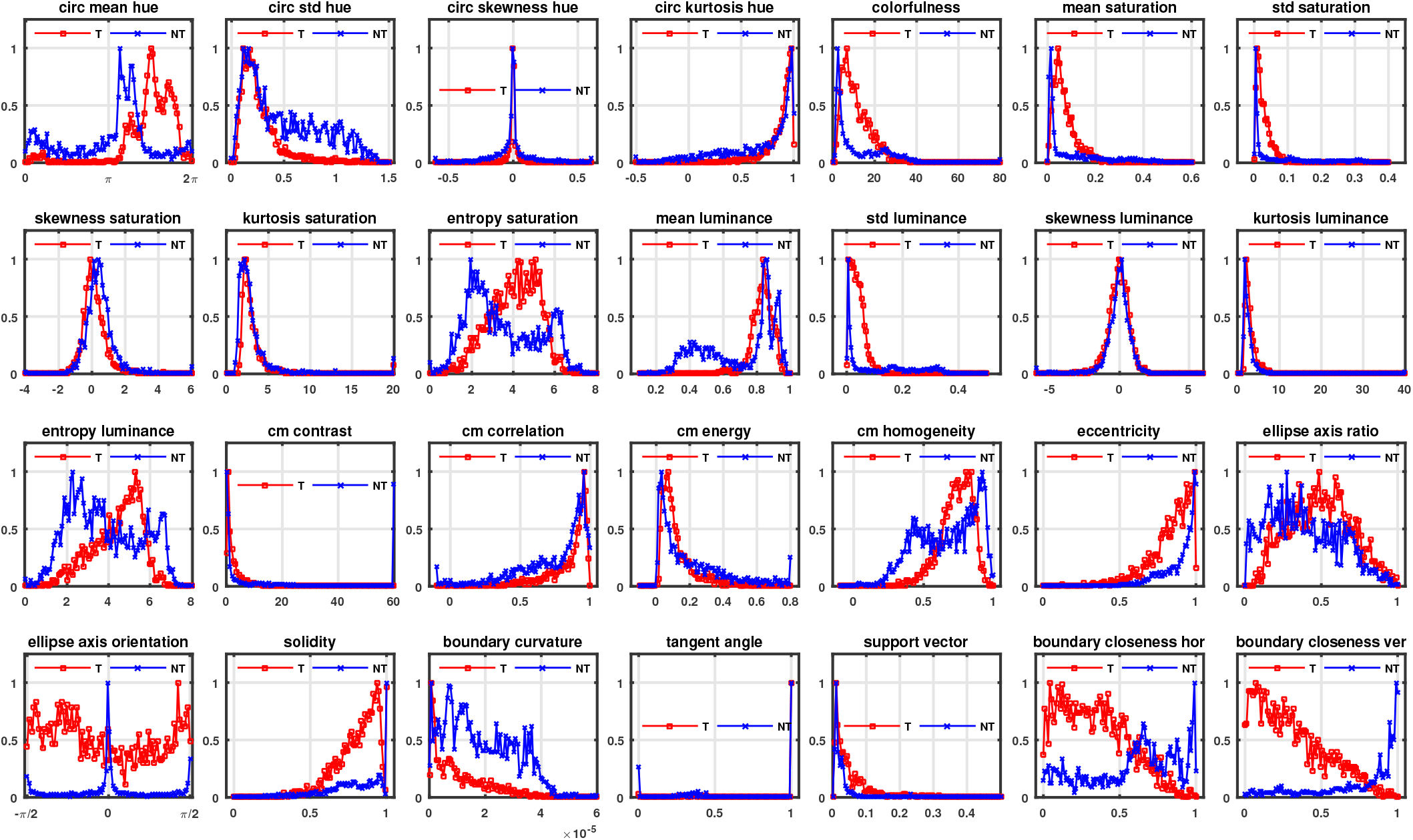
Distribution of individual image features for labelled tissue and non-tissue samples. The vertical axis indicates the normalized occurrences.

The class-relevance of individual features can be quantified using the Fisher score. The intuition behind it is that features with high separability should assign similar values to instances (samples) in the same class and different values to instances from different classes. The Fisher score for the i-th feature *F_i_* is computed by:

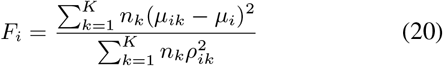

where *K* is the number of classes (*K* = 2 for binary classification), *n_k_* is the number of samples in the *k*-th class, *μ_i_* is the mean of the *i*-th feature, *μ_ik_* and *ρ_ik_* are the mean and the variance of the *i*-th feature in the *k*-th class, respectively. The numerator indicates the inter-class variance while the denominator indicates the sum of variance within each class. A larger Fisher score indicates that the feature is more discriminative.

The redundancy between image features *x_i_* and *x_j_* can be represented using the Pearson correlation coefficients *ρ_ij_*:

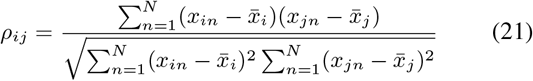

where *N* is the number of samples, 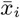 and 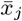 are the mean values of *i*-th and *j*-th features. Fig. 12 demonstrates the correlation matrices of features for labelled tissue and nontissue samples. These matrices contain the Pearson correlation coefficients between individual image feature and the others. Image features with significant linear dependency (i.e., the absolute value of the correlation is close to 1) can be reduced since they contain redundant information. The observation of correlation matrices reveals that there are similar linear relationship between features regardless of classes since the patterns of both matrices are largely similar. In addition, there is a strong negative linear relationship between the eccentricity and the ellipse axis ratio features, thus, eliminating either feature would not significantly influence the classification accuracy.

**Fig. 12.**
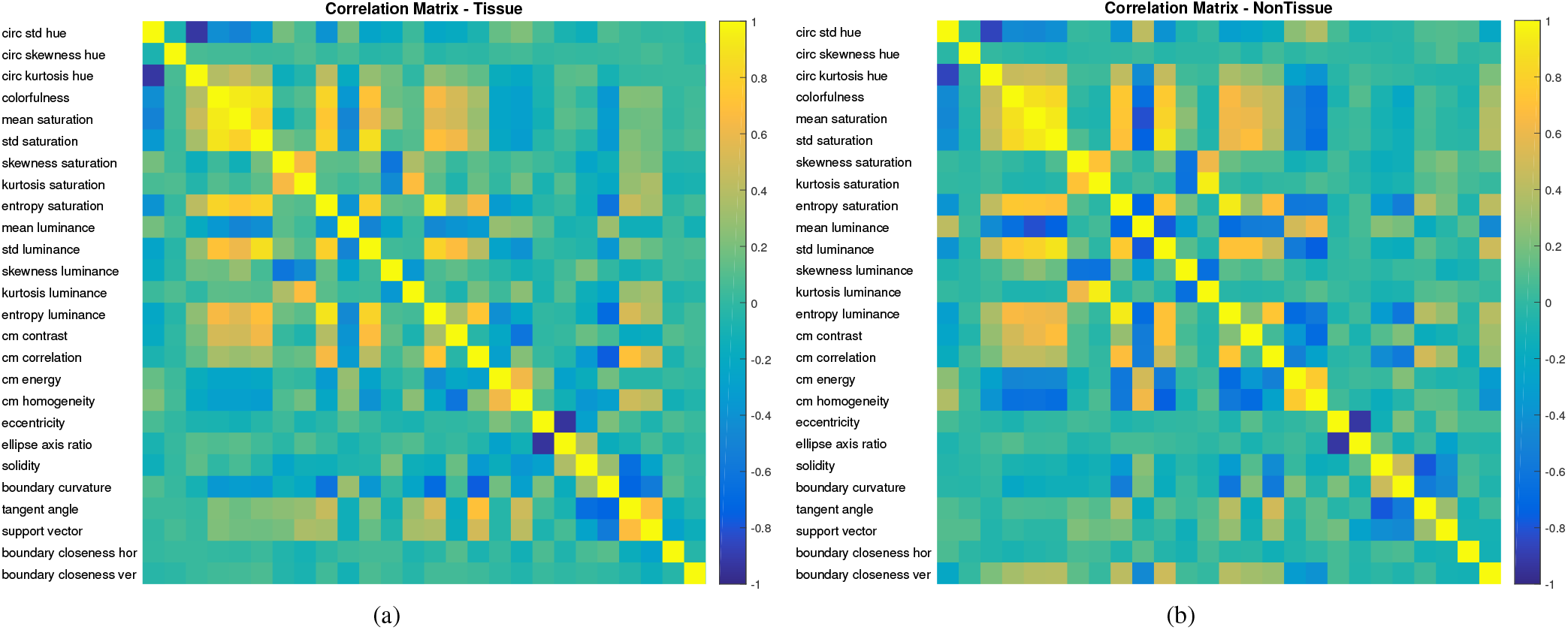
Correlation matrices for labelled tissue and non-tissue samples.

## V. Tissue Classification

In this work, we use the support Vector Machine (SVM) to classify tissue and non-tissue regions since it is a well-established machine learning solution for handling high dimensional data. The basic idea is to search for the best hyperplane in order to maximize the separation margin between two classes.

### A. Support Vector Machine

Consider we have *L* training samples, where each input x_*i*_ ∈ ℝ^D^ has *D* features (i.e., is of dimensionality *D*) and is in one of two classes *y_i_* ∈ {-1, +1}. The objective of SVM is to put a hyperplane in the middle of the two classes in a way that the distance to the nearest positive or negative example is maximized.

The SVM discriminant function has the form *f* (*x*) = *w^T^ x* + *b*, where *w* is the parameter vector and *b* is the bias or offset scalar. The classification rule is *sign*(*f*(*x*)) and the linear decision boundary is specified by *f* (*x*) = 0. If *f* separates the data, the geometric distance between a point *x* and the decision boundary is 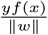.

Given training data {*x_i_,y_i_*}, the objective of SVM is to find a decision boundary *w* and *b* such that to maximize the geometric distance of the closer point, i.e..

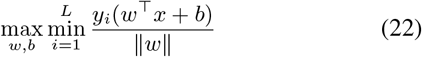

and it is equivalent to a quadratic programming problem as follows:

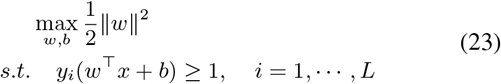

To handle linearly non-separable datasets, the constraints can be relaxed by making the inequalities easier to satisfy by introducing a slack variable *ξ_i_* ≥ 0 as follows:

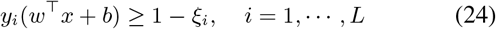

A point *x_i_* can satisfy the constraint (24) even if it is on the wrong side of the decision boundary. This problem can be restated as follows:

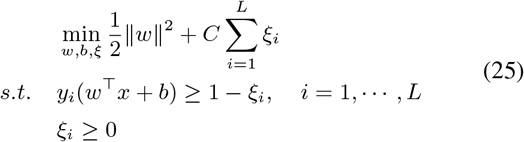

where *C* is the cost parameter for misclassification of training samples against simplicity of the decision boundary. A low *C* makes the decision surface smooth, while a high *C* aims at classifying all training samples correctly by giving the model freedom to select more samples as support vectors. The value of *C* can be set by cross validation.

The dual problem of (25) can be formulated by introducing Lagrange multipliers and it again becomes a quadratic programming problem as follows:

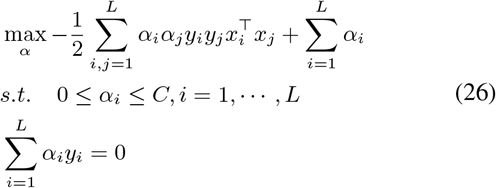

and the discriminant function is given by:

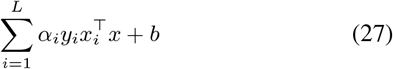

Since (26) and (27) only involve the dot product 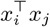 of examples, the kernel trick can be used. For kernel-base SVM, the dot product term 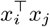 can be replaced with 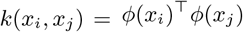. In this work, the kernel-based SVM is adopted since it allows for mapping of input data into high dimensional feature space so that nonlinear problem can be solved as a simpler linear classification problem. For kennel-based SVM, selecting an appropriate kernel function is of vital importance to ensure good performance. We exploit the Radial Basis Function (RBF) kernel *k*(*x_i_,x_j_*) = exp (-*γ*||*x_i_* – *x_j_*||^2^), since it requires less number of parameters to estimate than other alternatives, e.g., polynomial or sigmoid, while producing competitive performance in many scenarios. Intuitively, the *γ* parameter of RBF kernel defines how far the influence of a single training sample reaches, with low values meaning ‘far’ and high values meaning ‘close’. If *γ* is too large, the radius of the area of influence of the support vectors only includes the support vector itself and no amount of regularization with C will be able to prevent overfitting. When *γ* is very small, the model is too constrained and cannot capture the complexity or “shape” of the data.

Adopting the recommendation from [14], the optimal values of (*C, γ*) are identified for individual tests by performing a grid-search, i.e., examining various combinations of (*C, γ*) and choosing the one that yields the highest performance (i.e., high classification accuracy).

## VI. ROI

The ROI step is the final step in the pipeline. First, a bounding box is drawn around the tissue, this is the area that will be scaned at high magnification. A white balance box is placed on the background to colour correct the image. Finally, focus dots are placed on the tissue for high resolution scanning. Figure 13 shows a brightfield preview image with ROI and focus dots.

**Fig. 13.**
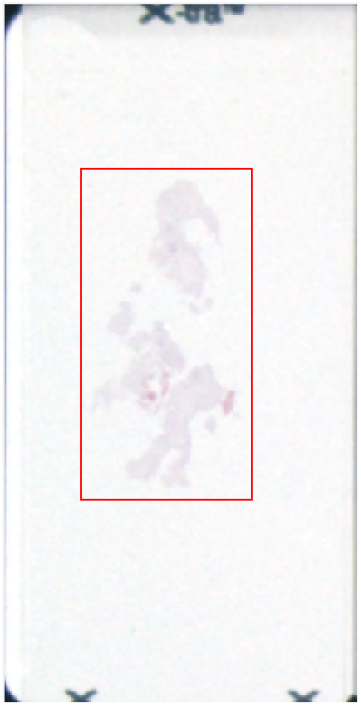
ROI applied to a preview image

## VII. Experimental Results and Discussion

### A. Database

The choice of appropriate image database is important for both training and validation of tissue classifier. With respect to tissue/non-tissue classification in whole slide imaging, there are lack of publicly available database. In this work, we have generated manually labelled database that allows for reliable training and validation of the proposed classifier. The labeling is done using semi-automated tagging tool. The slide images in the collected database contain tissue samples of diverse characteristics, i.e., varying tissue types, stain colors, sizes and distribution density with varying imaging conditions, i.e., acquisition devices and artifacts (e.g., color casting and faint tissues).

The manual labelling process initially applies a rough automated segmentation on slide images. It is followed by manual labelling performed by human operators that individual connected regions are manually labelled as ‘1’ or ‘0’ for tissue and non-tissue, respectively. For tissues with conflict labelling, a voting process is adopted. The labelled sets are validated by semi-experts, resulted in minor corrections. Finally, the resultant binary tissue map that contains regions of manually labelled tissue is collected as groundtruth. This process is done for 394 tissue images. The collected dataset is composed of three subsets (Fig 14): i) Mixed Preview Set: containing preview images with white to grayish background, high to medium quality tissues; ii) Tissue Snap Set: containing preview images with reddish color casting issues and blurred tissues; and iii) Set 1: containing preview images with bright white background and high quality crisp tissues. Detailed descriptions of individual subsets are demonstrated in Table I.

**Fig. 14.**
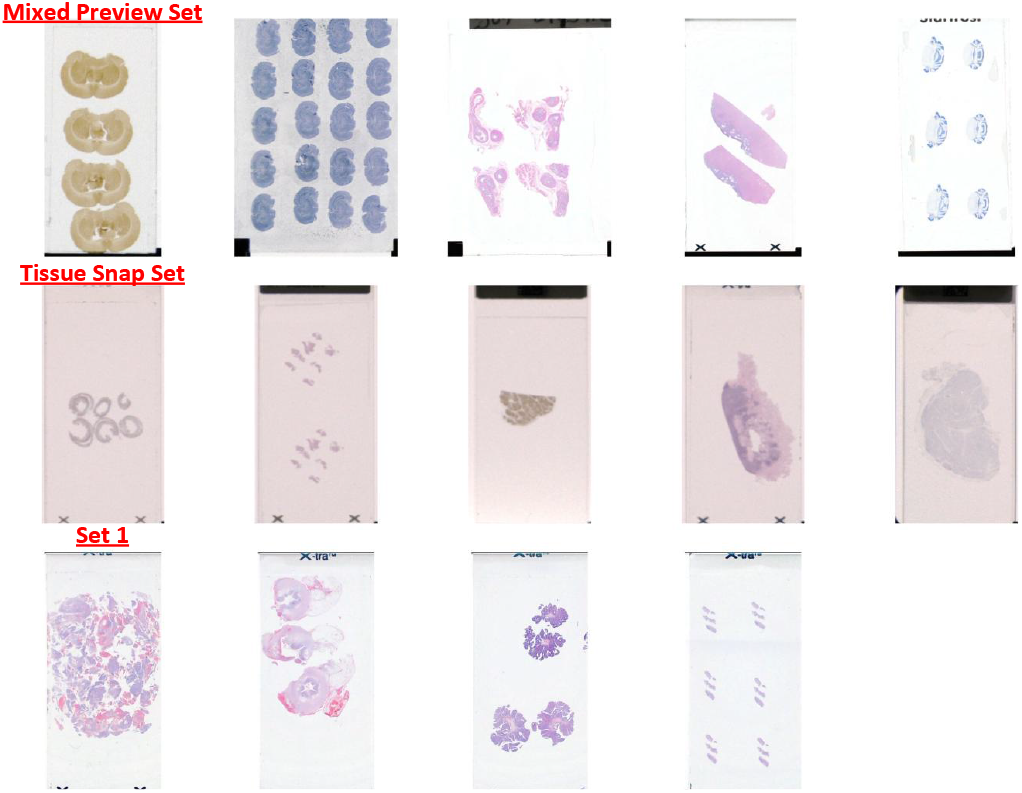
Tissue slide samples in the labelled datasets.

### B. Training Protocol

We incorporate the labelled tissue and non-tissue samples to evaluate the classification performance of the proposed classifier. Since the proposed approach requires a training process to calibrate a SVM classifier, we divide validation database into two randomly chosen non-overlapping subsets. Consider we have *N* = *N*_1_ + *N*_2_ labelled samples (*N*_1_ and *N*_2_ denote the number of tissue and non-tissue samples, respectively), where each input *x_i_* ∈ ℝ^D^ has *D* features and is in one of two classes *y_i_* ∈ {*ω*_1_, *ω*_2_} (*ω*_1_ and *ω*_2_ represent the tissue and nontissue class, respectively). If not explicitly stated otherwise, we divide the labelled samples into two randomly chosen non-overlapping subsets: 70% training and 30% testing for both tissue and non-tissue samples (i.e., training data X_*R*_ contains 0.7*N*_1_ tissue and 0.7 *N*_2_ non-tissue samples, whereas testing data X_E_ contains 0.3*N*_1_ tissue and 0.З*N*_2_ non-tissue samples). ^2^ To ensure that the results generalize across different training/test sets, we repeat this test procedure 200 times and the medians of classification accuracies are reported.

### C. Performance with Full Feature Set

Fig. 17 demonstrates the performance of the SVM-based classifier with varying ratio of training and testing sets. For this experiment, entire 28 image features are exploited for evaluation. For individual case, classification performance is reported with the optimal SVM parameters (*C,γ*), which are obtained from a gridsearch.

**Fig. 15.**
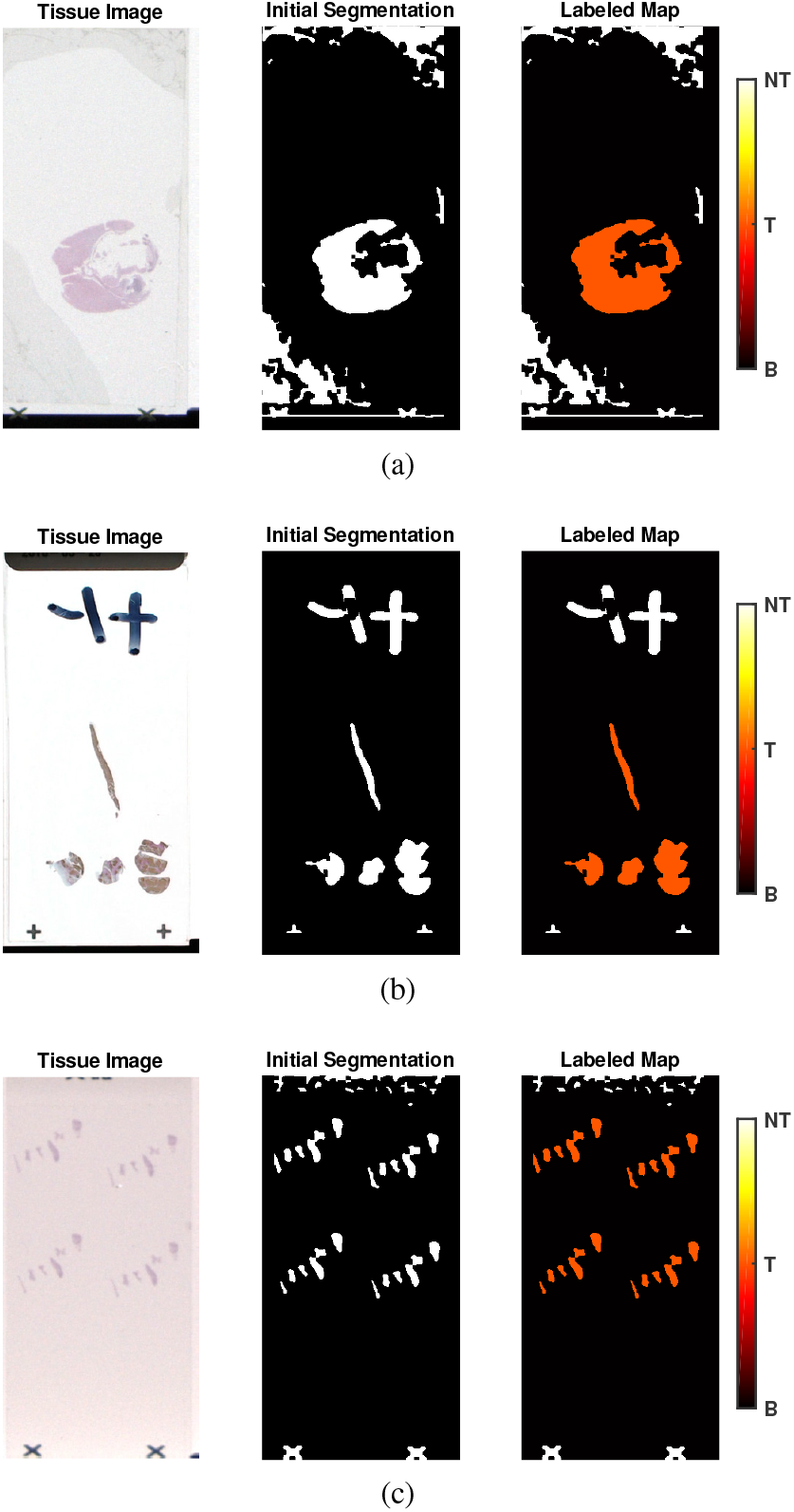
Visual examples of labelled tissue samples. The orange patches indicate the regions labelled as tissue (‘T’), whereas the white patches indicate the regions labelled as non-tissue (‘NT’).

**Fig. 16.**
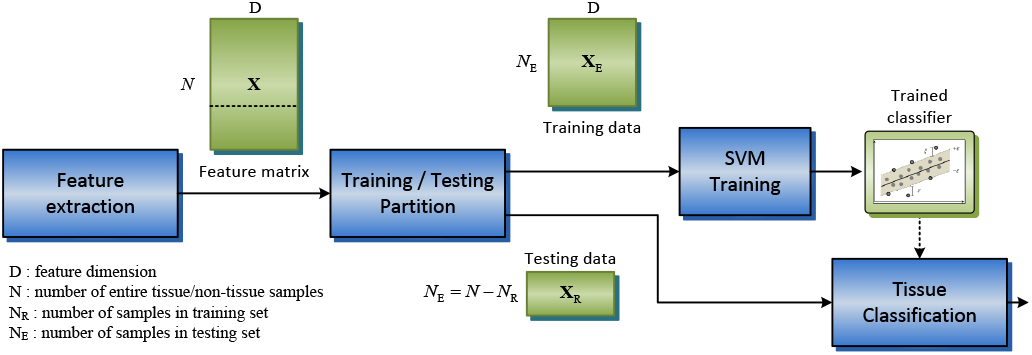
Workflow of training and testing data partition for the proposed tissue classifier.

**Fig. 17.**
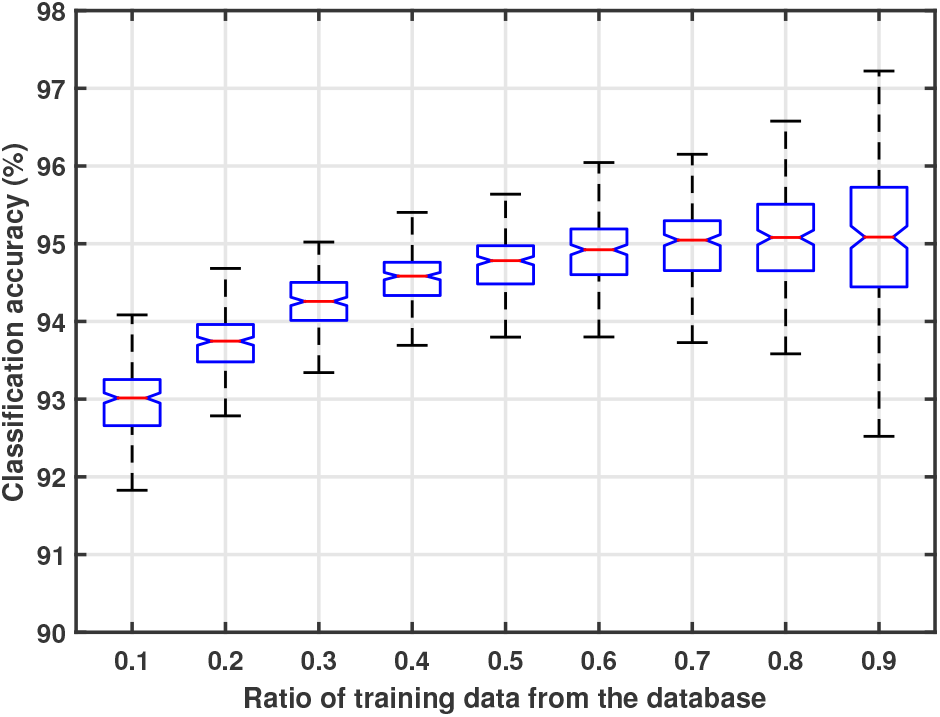
Performance of the SVM-based tissue classifier for different ratio of train/test split for validation database. The bottom and top edges of the blue box indicate the first quartile (Q1) and third quartile (Q3), respectively. The red line inside the blue box is the median (Q2) classification accuracy of 200 trials. The bottom and top ends of the whiskers represent the lowest accuracy within 1.5 interquartile range (IQR: distance between Q1 and Q3) of the Q1 and the highest accuracy within 1.5 IQR of the Q3. The notch extremes correspond to 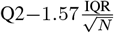 and 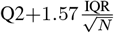, where *N* is the number of trials.

Fig. 17 shows that the median classification accuracy consistently increase as we increase the size of training data, indicating more stable performance of the classifier. Using more than 70% of data as training data yields statistically indistinguishable gain in terms of classification accuracy with 95% confidence level (since the notches in the boxplot overlap), indicating that we have sufficient amount of labelled data for training the SVM classifier. With large training size (i.e., 90%/10% train/test partition), the variances of reported classification accuracy are substantially larger than other cases since the size of testing data is relatively small. It demonstrates that the train/test split of 70%/30% is reasonable partition size for the given database to validate the tissue classifier.

Fig. 18 presents the gridsearch results of varying SVM parameters for three different partition ratio. It shows that the classifier performance for varying parameter remains unchanged over partition ratios, and the optimal performance can be achieved when the SVM parameters are configured as (*C,γ*) = (2^11^,2^-11^).

**Fig. 18.**
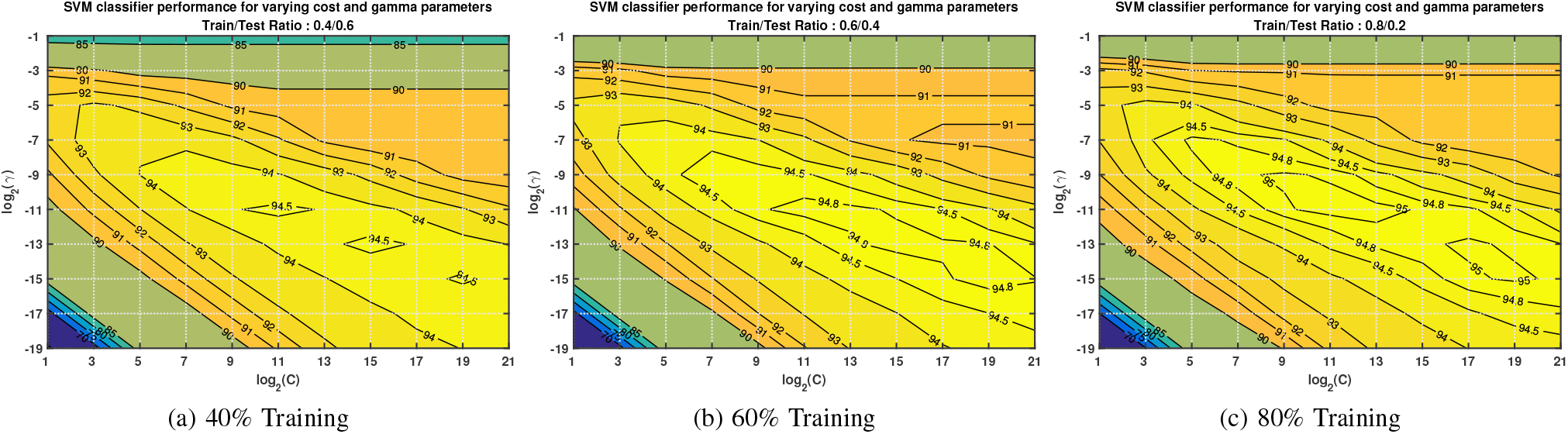
The gridsearch results of the SVM classifier for varying combinations of *γ* and *C* SVM parameters over three different train/test partition sizes.

### D. Performance with Reduced Feature Sets

High dimensional data could contain irrelevant and redundant information, which may lead the performance of learning algorithms to deteriorate. In this experiment, we evaluate the performance of classifier with existing feature extraction^3^ and feature selection techniques. Feature selection is the process commonly used for removing irrelevant and redundant features while maintaining acceptable classification accuracy. It should be distinguished from feature extraction that feature selection exploits a subset of the features, whereas feature extraction creates new features from functions of the original features.

#### 1) PCA-based Feature Extraction

Principal Component Analysis (PCA) is a non-parametric approach widely used for dimension reduction. Given a set of data on *D* dimensions, PCA aims to find a linear subspace of dimension *D*’ < *D* such that the data points lie mainly on this linear subspace. Such a reduced subspace attempts to maintain most of the variability of the original data. The steps for performing PCA in training and testing are summarized in the pseudo code shown in Fig. 19. The classifier is trained with the reduced data 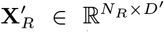 and tested on the reduced test data 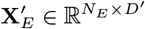, where *D*’ ≤ *D* denotes the number of reduced features. Fig. 20 shows the performance of the classifier for varying feature dimensions. The classifier maintains stable performance when feature dimension is set to *D*’ ≥ 15.

**Fig. 19.**
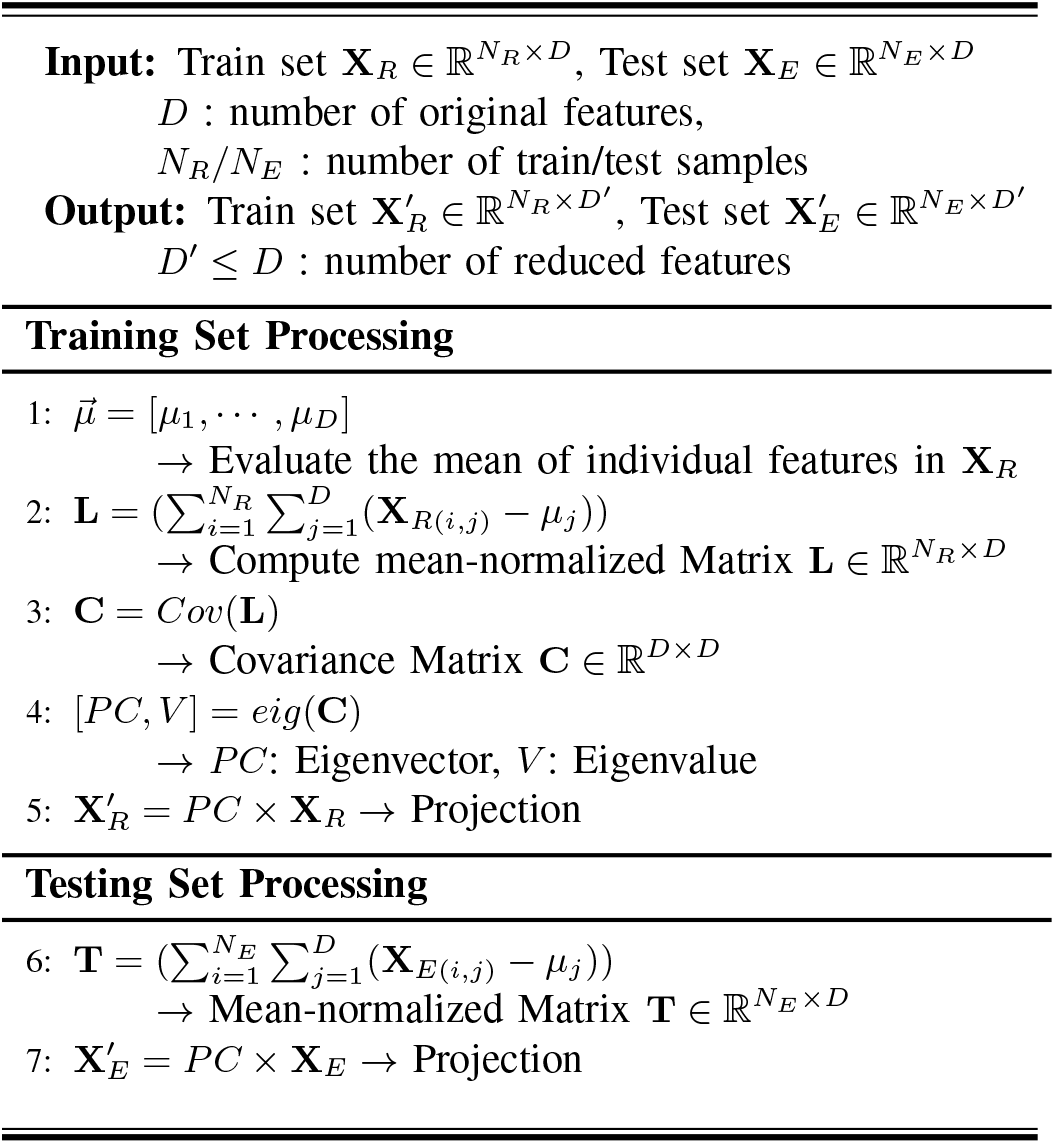
PCA-based feature reduction on training and testing data

**Fig. 20.**
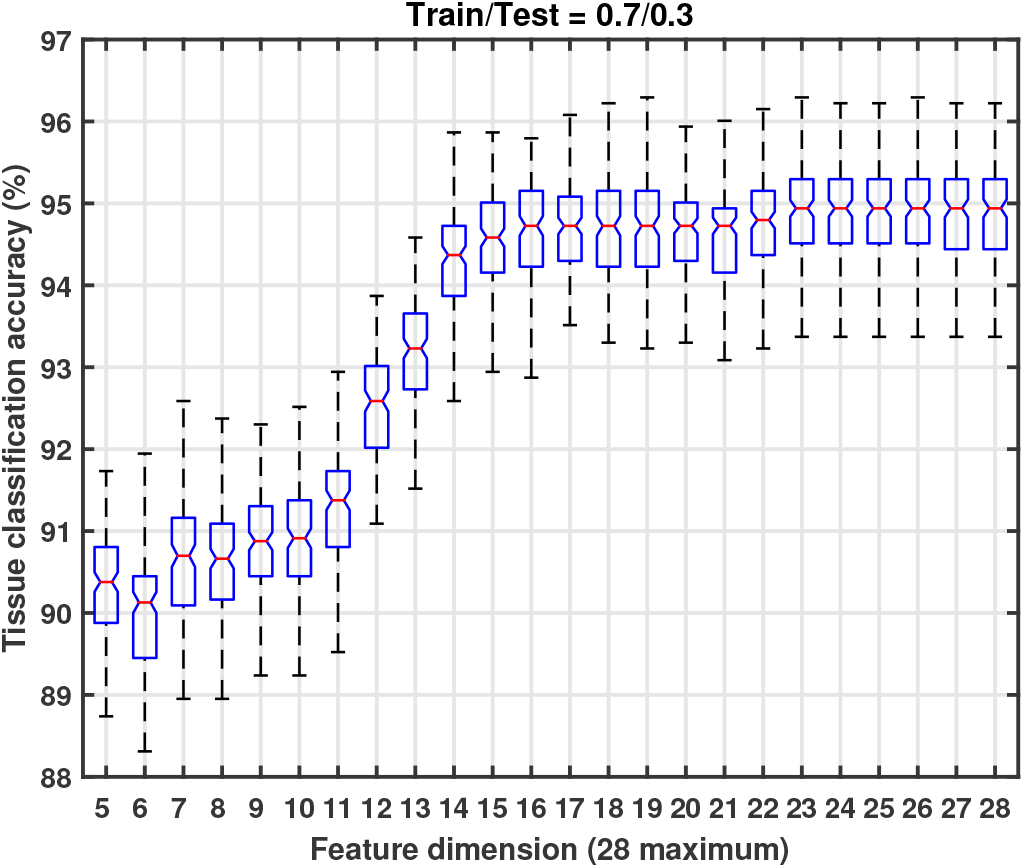
Performance of the tissue classifier for varying number of features with PCA dimension reduction.

#### 2) Feature Selection

In practice, identifying a subset of image features from entire 28 feature set is more efficient than the aforementioned PCA-based feature reduction technique which involves additional steps. In this section, entire 28 features are ranked based on feature ranking algorithms and the classifier performance on the feature subsets are examined as illustrated in Fig. 21.

**Fig. 21.**
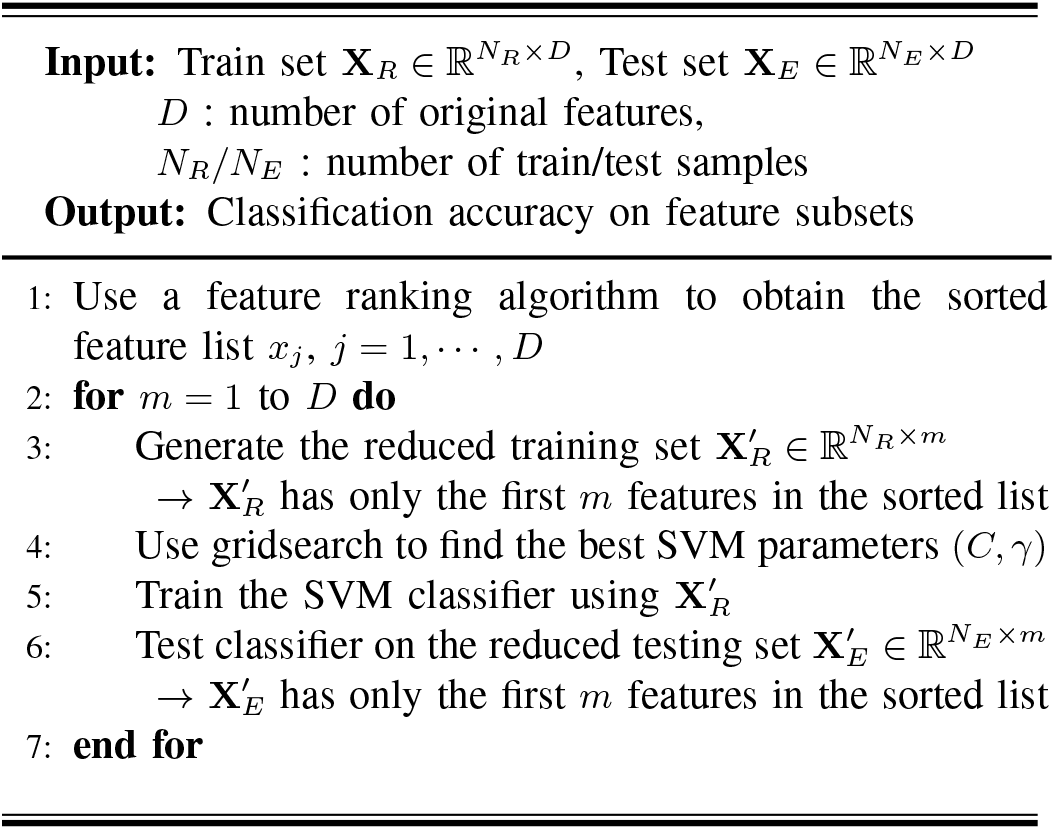
SVM-based classification with feature selection

The first exploited ranking algorithm is the Fisher score in (20), where the feature with lowest Fisher score has the least discriminative power. Fig. 22 illustrates the classification accuracy as a function of number of remaining features, indicating that the maximum accuracy can be achieved by eliminating four features with lowest Fisher score, i.e., the circular skewness of hue (f03), saturation kurtosis (f09), luminance skewness (f13) and luminance kurtosis (f14). It is notable that retaining only 15 features based on the Fisher score offers statistically indistinguishable classification accuracy compared to the maximum accuracy achievable with 24 features. It clearly indicates that there are some irrelevant features and the trained classifier performs more accurately by removing such features.

**Fig. 22.**
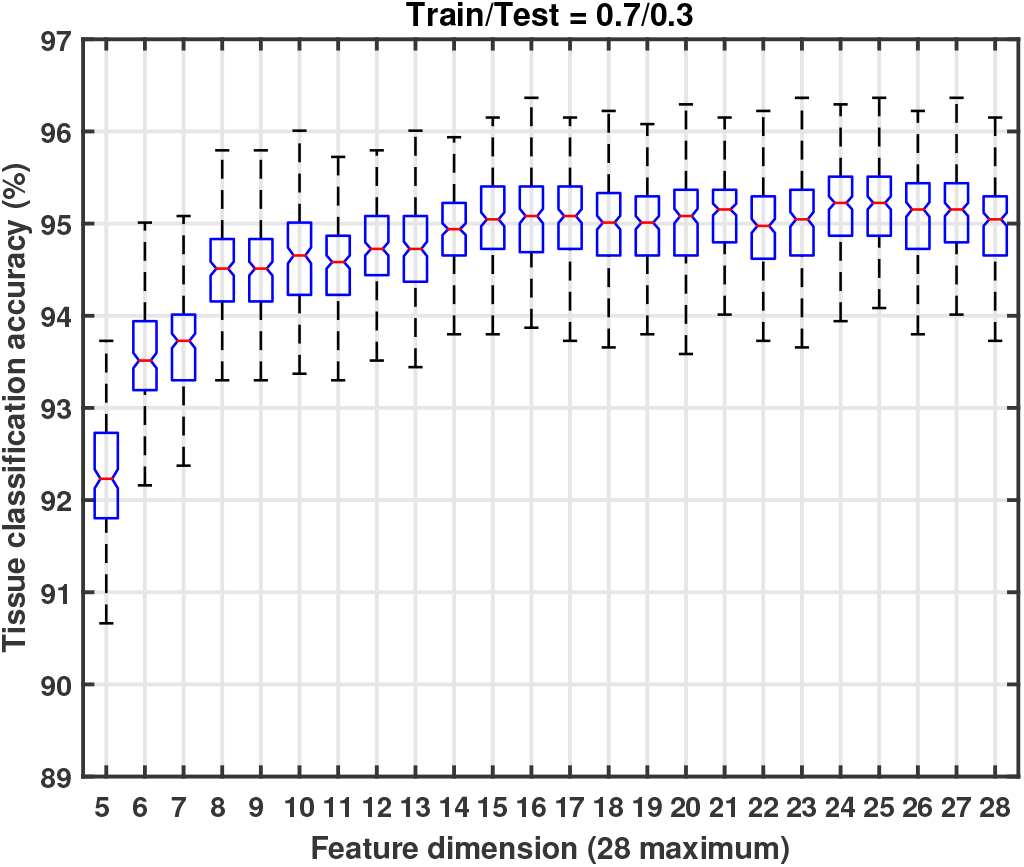
Performance of the tissue classifier for varying number of features selected using the Fisher score.

However, the Fisher score-based selection does not take into account the redundancy between image features. Therefore, we also exploit the minimum Redundance Maximum Relevance (mRMR) criterion [15] for feature selection which considers the tradeoff between relevance and redundancy. Consider the purpose of feature selection is to find a feature set *S* with *m* features {*x_i_*}. The maximum relevance condition aims to search features satisfying [15]:

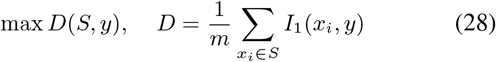

where *I*_1_(·) measures the relevance of feature *x_i_* to the class label *y*. This condition is to maximize the mean relevance value between individual feature *x_i_* and class *y*. Since it is likely that features selected according to the maximum relevance condition have dependency, the minimum redundancy condition selects mutually exclusive features [15]:

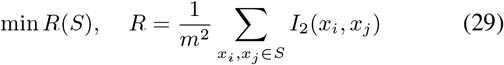

where *I*_2_(·) measures the redundancy between two individual features.

The mRMR criterion combines them into a single criterion by maximizing the difference between the relevance and redundancy conditions, i.e., max(*D – R*). The incremental search algorithm can be used to find the near-optimal solution for the mRMR criterion. Assuming that we have the feature set *S_m_* with *m* features, the algorithm aims to select the (*m* + 1) th feature from the set {Ω – *S_m_*} (all features except those already selected) by optimizing the following condition [15]:

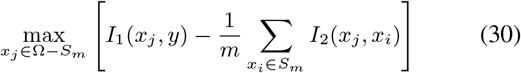

Another variant of the mRMR criterion is to maximize the divisive combination (quotient) of two conditions [16]. The optimization condition for the divisive combination becomes:

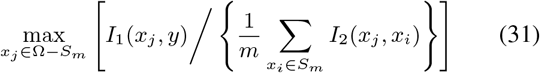

The appropriate selection of the measures *I*_1_(·) and *I*_2_(·) has a critical significance in the mRMR criterion-based feature selection. In its original form, the authors offer two options [16]: i) mutual information measure is used for both *I*_1_(·) and *I*_2_(·); and ii) the Fisher score and the absolute of Pearson correlation coefficient are used as *I*_1_(·) and *I*_2_(·), respectively.

Fig. 23 illustrates the classification accuracy as a function of number of remaining features with the mRMR-based feature selection. It shows that the best selection scheme for the given dataset is the divisive criterion with Fisher-Correlation measures. Another interesting observation is that the Fisher score-based selection produces slightly better result than the mRMR criterion even though it relevance and redundancy for feature selection.

**Fig. 23.**
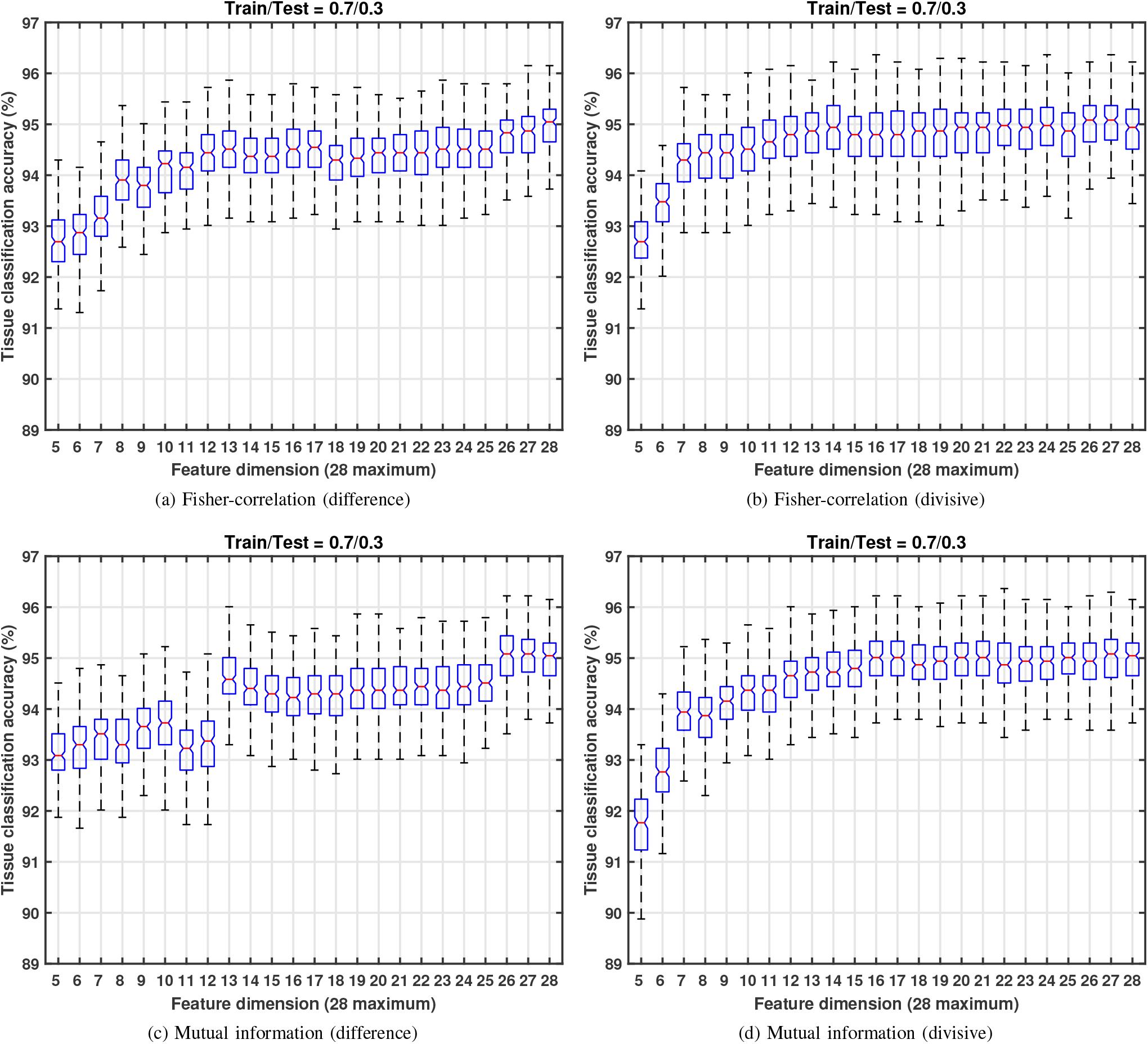
Performance of the tissue classifier for varying number of features with the mRMR criterion

## VIII. Darkfield Tissue Finding Package

### A. Differences That Arise With Darkfield

Darkfield images are different to brightfield images in three major ways.

- Brightfield images have a bright nearly white background on which darker regions of tissue are visible. Darkfield images are the opposite, the background is black and tissues appear as faint brighter regions.
- Brightfield images show tissues as a solid mass, whereas a darkfield image highlights the edges and vesicle of the tissue.
- The provided darkfield images are captured in grayscale whereas brightfield images where captured in three channel RGB.

Figure 24 shows a side by side comparison between a brightfield and a darkfield image of the same tissue sample.

**Fig. 24.**
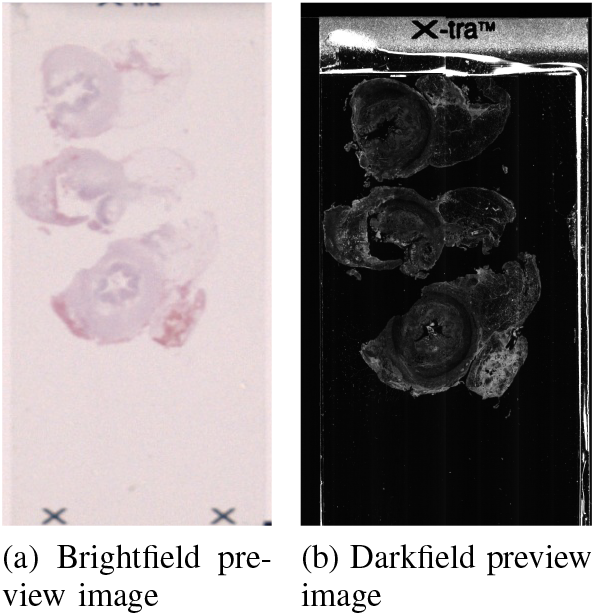
Darkfield preview image compared to a brightfield preivew image of the same tissue slide.

### B. Modifying Input Image

As previously mentioned, darkfield images have an inverse colour scheme to brightfield images. The background appears black instead of white and the tissue appears white rather than dark. To minimize the amount of changes that must be made to the brightfield tissue finding package it is desirable to make darkfield image resemble brightfield images. The darkfield tissue finding package therefore inverts the colour scheme of the inputed images. This is the first manipulation done to the image after it is loaded from memory. The MATLAB implementation of the code uses a MATLAB’s imcomplement function, while in C++ the original input image is subtracted from 255. The original input image is shown in comparison to the complemented image in figure 25.

**Fig. 25.**
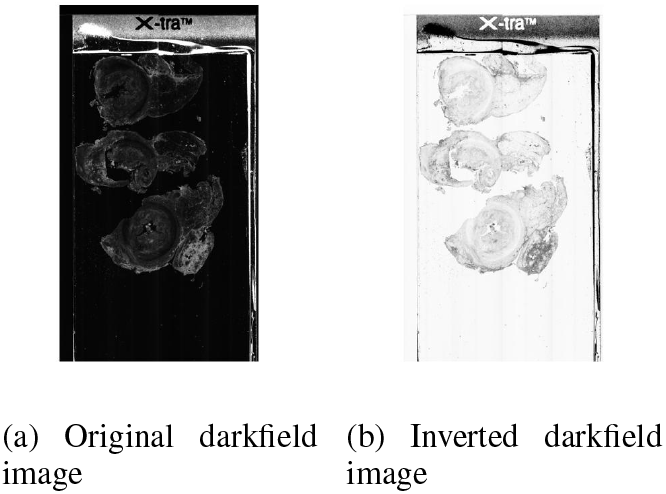
Colour of the input image is inverted. This is the first step in the darkfield tissue finding package. It is done so the input image resembles brightfield images.

### C. Increasing Sensitivity Of Mask

Because darkfield imaging only captures scattered light which is significantly dimmer than the direct light captured by brightfield imaging, the tissues seen on a darkfield image are significantly dimmer than one seen on a brightfield image. To compensate for this the darkfield tissue finding package must be made more sensitive.

As previously mentioned, several masks are created throughout the tissue segmentation pipeline. For the darkfield tissue segmentation package the RGB mask and L mask were made more sensitive. This was done by multiplying the threshold value produced by the existing adaptive threshold function by 1.205 and 1.009 for the RGB and L mask respectively. Multiplying the adaptive threshold value by a constant is a simple solution that allows brightfield code to be reused.

In brightfield, a resultant mask is made from combining an RGB mask and an L mask using an AND function. In darkfield, however, this leads to undermasking. It was found that in darkfield, the RGB mask was more effective at capturing the tissue body, whereas the L channel mask was more affective at capturing the tissue edges and vesicles. Therefore, in darkfield the resultant mask is created by combing the RGB and L masks using an OR function. The mask outputs are seeen in figure 26

**Fig. 26.**
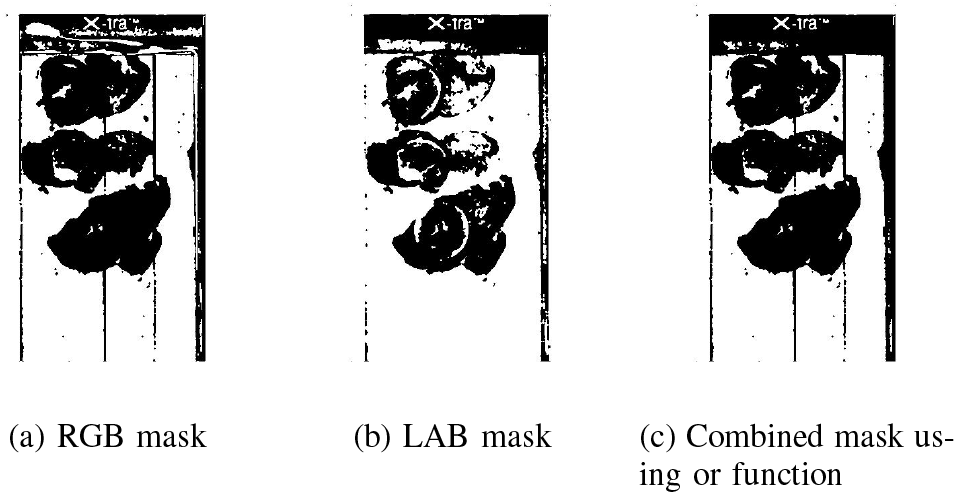
Darkfield mask generation. RGB and L channel mask sensitivity has been increased. The masks are now combined using an OR function rather than an AND function to retain more detail.

### D. Boundary Detection

In darkfield images the slide’s edges are very prominent. If a tissue is placed near these edges it becomes very difficult to classify and segment the image. Edge detection is done in brightfield using first and second derivatives but due to the noisy and veccile nature of darkfield images, this method is not affective in darkfield. A new mask based boundary detection method was added to the darkfield tissue findining pipeline.

The new boundary detection method uses the resultant mask from RGB and L channel thresholding to determine the edges of the slide. This step therefore, proceeds the resultant mask generation step. It precedes any resultant mask enhancement and eliminate tensor feature steps.

The slide’s edge artifacts along the horizontal axis are found using column wise summation. The slide’s edge artifacts along the vertical axis are found using row wise summation. Because the artifact have a straight either vertical or horizontal nature they apprear as discontinuities in the row wise and column wise summation seen in figure 27. Identifying these discontinuities reveals the location of these artifacts. The artifacts are cropped out and the mask is padded with zeros to its original size, seen if figure 28.

**Fig. 27.**
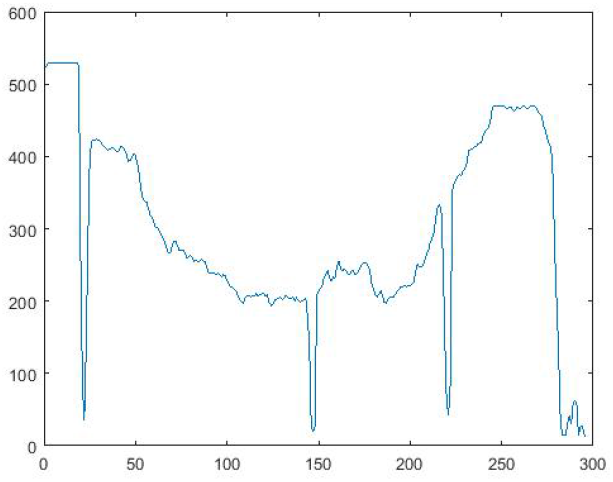
Plot of column-wise intensity summation. The slide artifacts cause the two discontinuities seen on the left and right sides of the curve. The dip in the center is caused by the tissue located there. The gradual change in values suggests the tissue has a spherical shape.

**Fig. 28.**
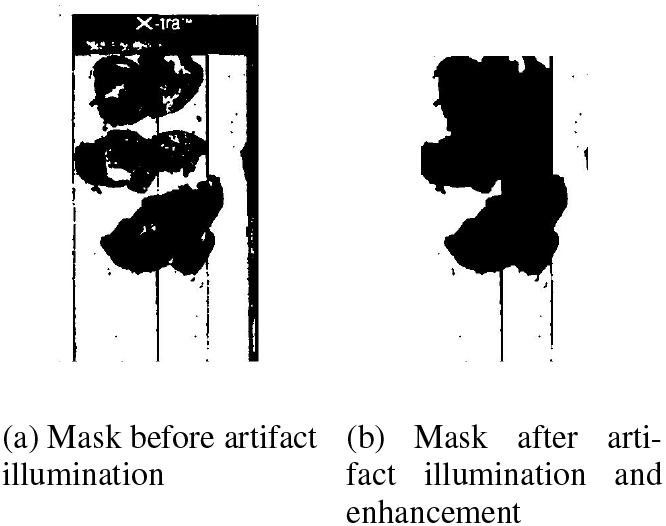
Boundary elimination. After the discontinuities in the row and column wise summation vectors are found, the slide artifacts are cropped out and padded with empty space.

### E. Classification

The classification step of the pipeline remains largely unchanged from its brightfield implementation. The only change done to this step is the removal of all colour features from the feature extraction step and retraining the SVM model to ignore said colour features. Features number 11,12, 15, 16, 17, 18, 20, 21, 22, 24, 25, 27, 28 from the table in figure **??** are used to classify darkfield preview images.

### F. Focus Dot Placement

Focus dots are needed to capture in focus high magnification images. At high magnification, any small change in the distance between the camera lens and the object can cause the image to go out of focus, become blurry and illegible. The distance between the camera and the tissue’s surface often varies because the thickness of the tissue varies. Huron’s tissue slide scanner is able to compensate for changes in the tissue’s thickness by moving the stage along the z i.e. vertical axis. Focus dots are locations for which the stage’s optimal z location that produces the most in focus image has been calculated. The values for these coordinates are then extrapolated and estimated for entire tissue region. The focus dot implementation uses a grid to segment the preview image into evenly spaced regions. The tissue area as a percentage of the entire area of each space is calculated. If said percentage is below a reject limit that space is ignored. If the percentage is above an acceptance limit a focus dot is placed in the middle of that area. Finally if the tissue area percentage is neither less than the reject limit nor greater than the accept limit the segment is further divided into four quadrants and the previous steps are repeated. Figure 29 shows a diagram of the focus dot placement algorithm and figure 30 shows focus dots placed on a preview image.

**Fig. 29.**
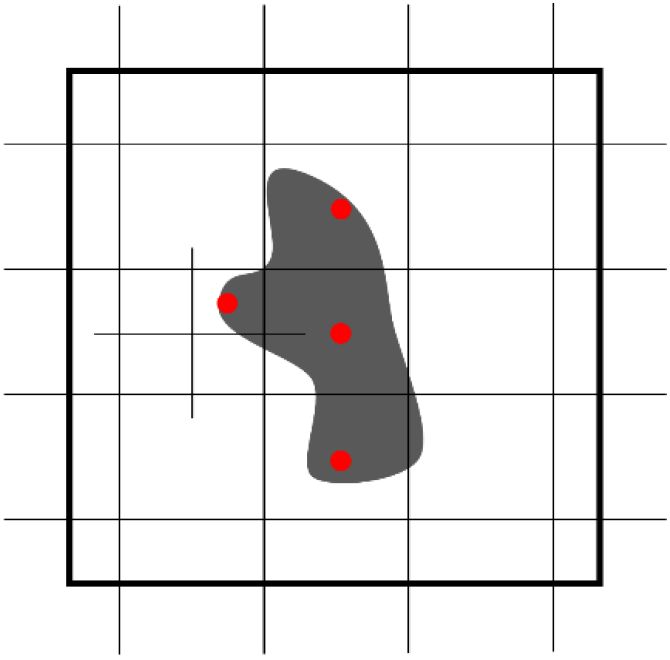
Diagram showing the location the grid based focus dot placing method. Initially a grid is placed above the preview image, dividing the image in to sections. In sections with a tissue area above a predefined threshold a focus dot is placed in the middle of the section. Sections with tissue area below a predefined threshold are ignored outright and no focus dot is placed there. Section with a tissue area that is between these two thresholds are divided into four smaller regions and the process is repeated.

**Fig. 30.**
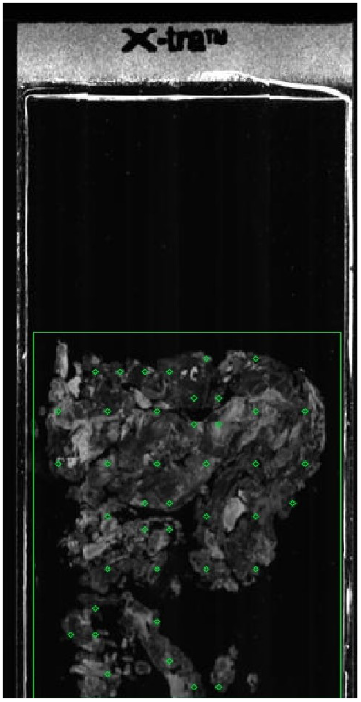
Figure showing focus dot placing algorithm in practice on a darkfield preview image

### G. Results for Darkfield Imaging

#### 1) Brightfield Stained Tissue Samples

The segmentation performance achieved with brightfield stained tissue samples satisfies the project requirements outlined above. The tissue segmentation package is able to consistently identify tissue samples and segment them from the background. The classification step is able to correctly classify tissue samples in the vast majority of cases, earring on the side of caution in difficult cases. Custom RoI placement gives the user more control of the classification process. By ignoring regions of artifacts, more accurate segmentation is achieved. Focus dots are placed on the tissue with holes and fat regions inside the tissue being avoided. The figures below demonstrate the segmentation results.

Some limitations include failing to classify out some artifacts leading to larger than required ROIs and failing to place focus dots throughout the entire region of tissue, perceiving instead dimmer regions of tissue as holes. Figure 31 shows segmentation results with automatic ROI, figure 32 shows segmentation results with custom ROI.

**Fig. 31.**
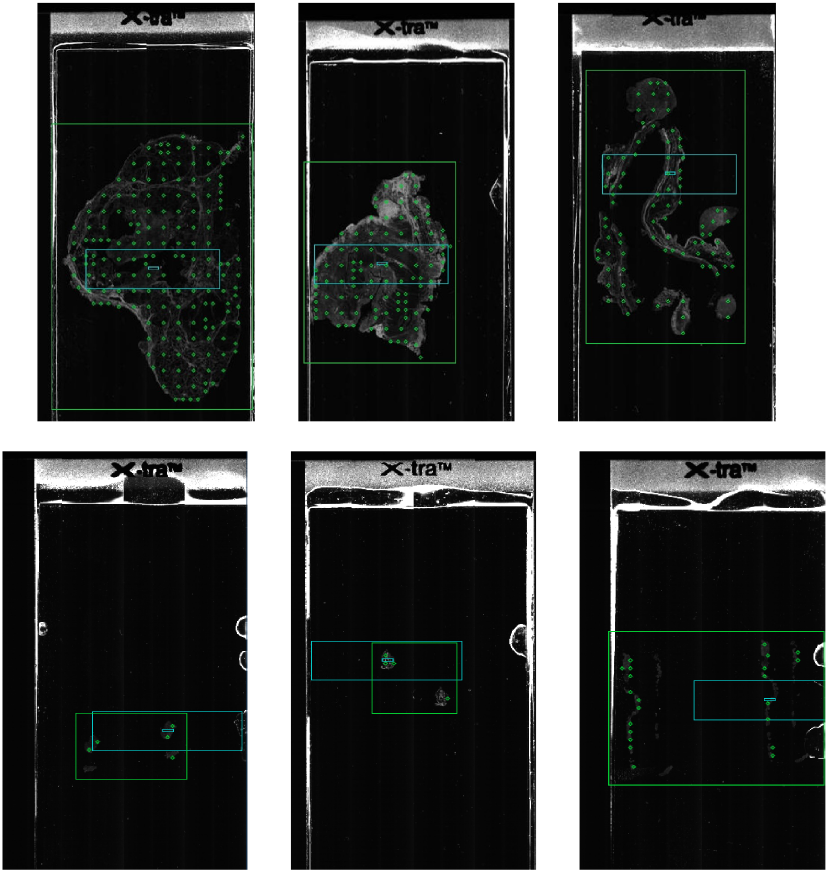
Automatic ROI results

**Fig. 32.**
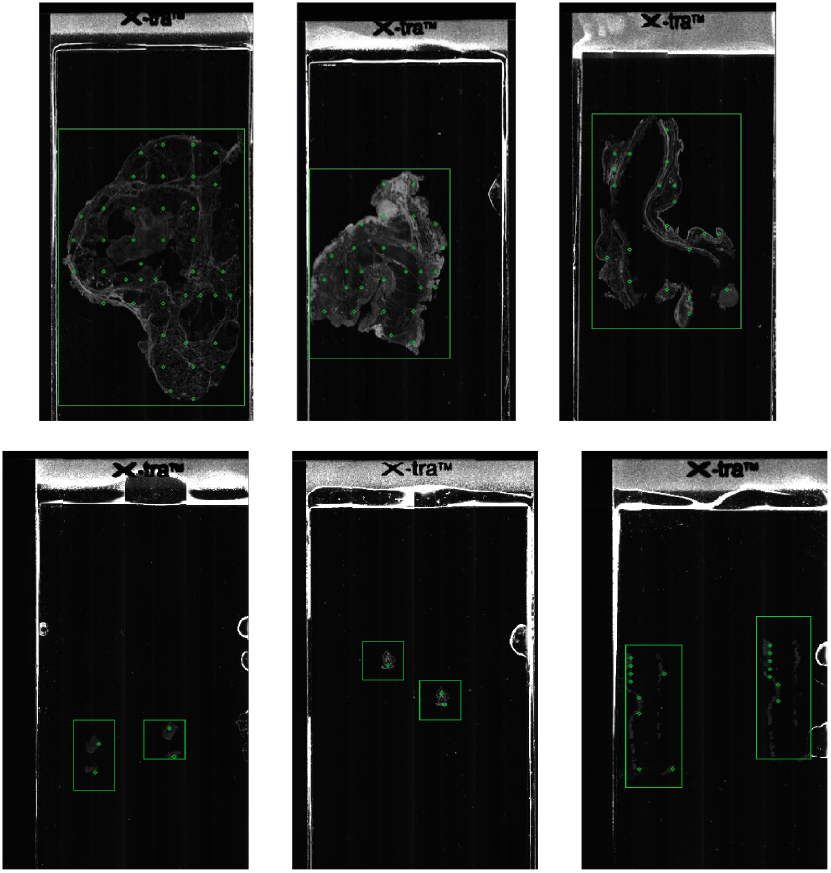
Custom ROI results

#### 2) Florescence Tissue Samples

Fluorescence tissue samples were significantly harder to segment for two reasons. Firstly, unlike the brightfield dye the fluorescence dye is clear making the tissue samples appear fainter on the darkfield preview images. secondly and more significantly the fluoresce tissue samples were quite badly scratched from a lifetime of use. The figure 33 demonstrate the original fluorescence preview image and an enhanced version to show the extent of the scratching and scuffing sustained by the tissue samples.

**Fig. 33.**
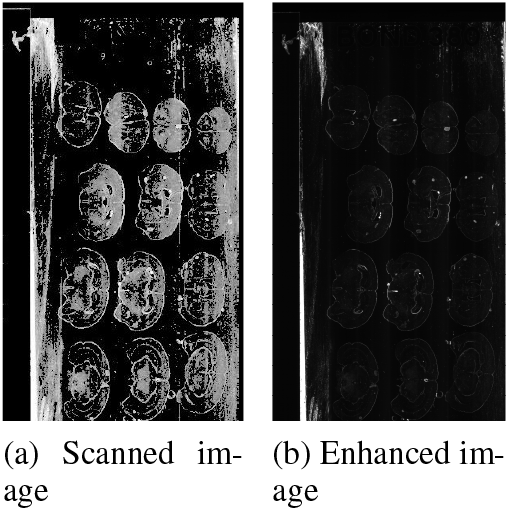
Figure showing the extent to which flouresense tissue samples are damaged. By changing the gamma level the scratches are highlighted.

The tissue segmentation package was unable to segment these preview images. It was unable to separate the tissue samples from the background and classify out the artifacts. The tissue segmentation output is shown in figure 34.

**Fig. 34.**
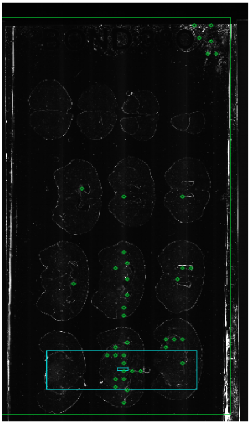
Tissue finding package results with florescence stained image

## IX. Software Integration

The proposed method was implemented in C++, OpenCV libraries and LIBSVM tools then integrated to Huron TissueScope Scanner to facilitate automated scan preparation. The preview image is acquired at 100 um resolution by both Logitech Webcam and Flir blackfly camera. Some of procedures including image segmentation are optionally adjusted by user-specified configuration such as crop size and masking. once ROIs are determined, a recursive depth-search is employed to prune or merge them depending on customized scanning mode (e.g., single ROI and multi ROIs). Focus dot placement is configured by adjusting grid size and target density. This method is also extended to execute in semi-automatic manner, which takes user-specified ROIs as an input to restrict tissue finding search and focus dot placement.

## Footnotes

1 This issue is often introduced because of morphological operations during segmentation task.

2 Please see Section VII-C for the reason for this partition ratio.

3 The purpose of this feature extraction is to remove data redundancies in the extracted image features. It should be distinguished from the image feature extraction process described in Section IV.

## References

[1] F. Ghaznavi et al., “Digital imaging in pathology: whole-slide imaging and beyond,” Annual Review of Pathology: Mechanisms of Disease, vol. 8, pp. 331–359, 2013.

[2] E. Goacher et al., “The diagnostic concordance of whole slide imaging and light microscopy: a systematic review,” Archives of pathology & laboratory medicine, vol. 141, no. 1, pp. 151–161, 2017.

[3] E. Abels and L. Pantanowitz, “Current state of the regulatory trajectory for whole slide imaging devices in the usa,” Journal of pathology informatics, vol. 8, 2017.

[4] S. Mukhopadhyay et al., “Whole slide imaging versus microscopy for primary diagnosis in surgical pathology: a multicenter blinded randomized noninferiority study of 1992 cases (pivotal study),” The American journal of surgical pathology, vol. 42, no. 1, p. 39, 2018.

[5] U. Food, D. Administration et al., “Technical performance assessment of digital pathology whole slide imaging devices. 2016; 81 fr 23306: 23306-23307,” 2018.

[6] A. J. Evans et al., “Us food and drug administration approval of whole slide imaging for primary diagnosis: A key milestone is reached and new questions are raised,” Archives of pathology & laboratory medicine, vol. 142, no. 11, pp. 1383–1387, 2018.

[7] A. F. Frangi et al., “Multiscale vessel enhancement filtering,” in International conference on medical image computing and computer-assisted intervention. Springer, 1998, pp. 130–137.

[8] M. S. Hosseini and K. N. Plataniotis, “Derivative kernels: Numerics and applications,” IEEE Transactions on Image Processing, vol. 26, no. 10, pp. 4596–4611, 2017.

[9] M. S. Hosseini and K. N. Plataniotis, “Finite differences in forward and inverse imaging problems:Maxpol design,” SIAM Journal on Imaging Sciences, vol. 10, no. 4, pp. 1963–1996, 2017.

[10] A. R. Smith, “Color Gamut Transform Pairs,” ACM SIGGRAPH Computer Graphics, 1978.

[11] A. Pewsey, “The large-sample joint distribution of key circular statistics,” Metrika, vol. 60, no. 1, pp. 25–32, 2004.

[12] D. Hasler and S. E. Suesstrunk, “Measuring colorfulness in natural images,” in Proc. SPIE, vol. 5007, 2003, pp. 87–95.

[13] W. Gomez, W. C. A. Pereira, and A. F. C. Infantosi, “Analysis of cooccurrence texture statistics as a function of gray-level quantization for classifying breast ultrasound,” IEEE Transactions on Medical Imaging, vol. 31, no. 10, pp. 1889–1899, Oct 2012.

[14] C.-W. Hsu, C.-C. Chang, and C.-J. Lin, “A practical guide to support vector classification,” Dept. Comput. Sci., National Taiwan Univ., Tech. Rep., 2003.

[15] H. Peng, F. Long, and C. Ding, “Feature selection based on mutual information criteria of max-dependency, max-relevance, and min-redundancy,” IEEE Transactions on Pattern Analysis and Machine Intelligence, vol. 27, no. 8, pp. 1226–1238, Aug 2005.

[16] C. Ding and H. Peng, “Minimum redundancy feature selection from microarray gene expression data,” Journal of Bioinformatics and Computational Biology, vol. 03, no. 02, pp. 185–205, 2005.

